# Insights into human outer kinetochore assembly and force transmission from a structure-function analysis of the KMN network

**DOI:** 10.1101/2023.08.07.552315

**Authors:** Soumitra Polley, Tobias Raisch, Marie Koerner, Melina Terbeck, Frauke Gräter, Stefan Raunser, Camilo Aponte-Santamaría, Ingrid R. Vetter, Andrea Musacchio

**Author notes:** **Equal contribution** (Soumitra Polley, Tobias Raisch).

## Abstract

The biorientation of chromosomes during cell division is necessary for precise dispatching of a mother cell’s chromosomes into its two daughters. Kinetochores, large layered structures built on specialized chromosome loci named centromeres, promote biorientation by binding and sensing spindle microtubules. The kinetochore outer layer consists of a 10-subunit apparatus comprising Knl1C, Mis12C, and Ndc80C subcomplexes (KMN network). The KMN network is highly elongated and docks on kinetochores and microtubules using interfaces at its opposite extremes. Here, we combine cryo-EM reconstructions and AlphaFold2 predictions to generate a model of the KMN network that reveals all intra-KMN interfaces. We identify and functionally validate two interaction interfaces that link Mis12C to Ndc80C and Knl1C. Through targeted interference experiments and molecular dynamics simulations we demonstrate this mutual organization stabilizes the KMN network. Our work reports the first comprehensive structural and functional analysis of the microtubule binding machinery of kinetochores and elucidates a path of microtubule-generated force transmission

## Main

With approximately thirty core subunits and myriads of regulatory and accessory subunits, kinetochores are among the largest and functionally most complex molecular machines in the cell ^1^Their fundamental function is to provide a self-perpetuating platform for efficient capture of spindle microtubules during cell division. In most species, kinetochore composition and organization are broadly conserved ^2,3^. In their simplest form, identified in *Saccharomyces cerevisiae* and other yeast species, kinetochores assemble as a single chromatin-anchored complex that captures a single microtubule ^4^. In most other species, this “point” configuration is implemented in a more complex “regional” version that captures multiple microtubules and that is probably generated by convoluting a “point” structure with multiple adjacent docking sites on a specialized confined chromatin domain, the centromere ^1,4^. The latter may reside at the end of the chromosome (telocentric chromosome), in the middle (metacentric), or somewhere in between (acrocentric and sub-metacentric). In yet another configuration, named holocentric, multiple kinetochore-docking sites on chromatin are distributed along the entire length of the chromosome^5^.

At low resolution, kinetochores appear as layered structures, with a centromere proximal layer, the inner kinetochore, and a centromere distal layer, the outer kinetochore ^1^. The inner layer incorporates the subunits of the so-called constitute centromere-associated network (CCAN, also known as Ctf19 complex in *S. cerevisiae*). The outer layer, whose assembly requires specialized sites on CCAN, consists of the 10-subunit KMN network super-assembly ^6,7^. KMN network subunits are in turn distributed into three subcomplexes (Figure 1a), named the Knl1 complex (Knl1C, consisting of KNL1 and ZWINT), the Mis12 complex (Mis12C, consisting of DSN1, MIS12, NSL1, and PMF1 subunits), and the Ndc80 complex (Ndc80C, consisting of NDC80/HEC1, NUF2, SPC24, and SPC25) ^8^. During mitosis, the KMN becomes recruited to the CCAN through Mis12C, which interacts directly with CENP-C and CENP-T, two CCAN subunits ^9-18^. Both interactions are enhanced by phosphorylation of the Mis12C DSN1 subunits on conserved residues (Ser100 and Ser109 in humans) ^14,19-24^. Responsible for these phosphorylation events that explain the stabilization of outer kinetochore assembly during mitosis is the prominent mitotic kinase Aurora B ^19,24-26^.

**Figure 1.**
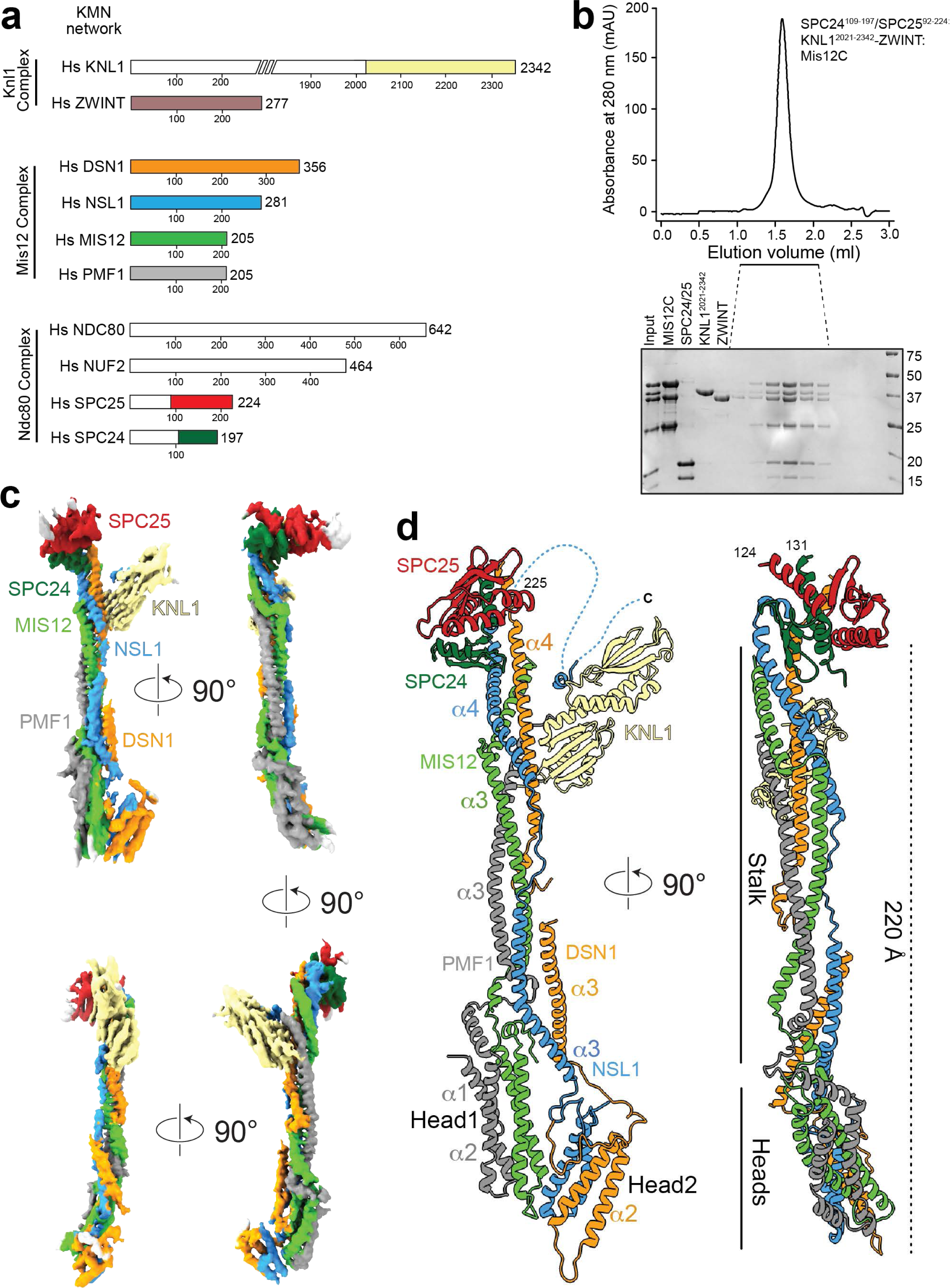
Structural organization of the KMN network. (**a**) Schematic organization of the KMN subcomplexes. Fragments included in reconstitution for cryo-EM analysis are shown with different colors, omitted fragments are in white. (**b**) Size-exclusion chromatography of the reconstituted KMN assembly on a Superdex S200 5/150 column. (**c**) Four rotated views of the experimental map. (**d**) Cryo-EM model of the structured core of the KMN network in cartoon representation.

Within the KMN, Mis12C has emerged as an interaction hub that coordinates Ndc80C and Knl1C ^27,28^. Specifically, Mis12C recruits Ndc80C through interactions with the RWD (C-terminal RING finger, WD repeat, DEAD-like helicases) domains of the SPC24/SPC25 subcomplex, and Knl1C through interactions with the C-terminal RWD domain of KNL1 ^12,13,15,29^. High-resolution structures of the Mis12C, of the KNL1 tandem RWD domains that interact with Mis12C, and of various segments of the Ndc80C, including in complex with microtubules, have been reported ^12,15,22,29-37^. The Mis12C forms a moderately elongated (∼22 nm) cylindrical structure with two head domains at the CCAN-binding end ^22,33^. Each of the four subunits of Mis12C is helical and runs in a parallel array along most of the long axis of the complex, with all N-termini at one end and all C-termini at the other. A low-resolution negative-stain electron microscopy (EM) analysis demonstrated that the KNL1 RWD domain docks on the side of the Mis12 cylinder, but the details of this interaction and the orientation of the tandem RWD domains in this arrangement remain obscure ^29^.

Low-resolution rotary-shadowing analyses on Ndc80C revealed an even more elongated structure (∼65 nm) with an overall shaft shape showcasing globular domains at each end ^13,38,39^. These consist of microtubule-binding calponin-homology domains at the microtubule-associated end, and – as already indicated – Mis12C-binding RWD domain at the centromere-targeting end. When combined, the Mis12C and Ndc80C add their length in series, generating “antennas” with a combined length of almost 90 nm ^13^.

Despite this progress, several crucial aspects of the organization of the KMN network remain unexplained. Specifically, the interfaces that allow Ndc80C and Knl1C to interact with the Mis12C hub are not understood on a structural level. Both interactions engage RWD domains, but there appears to be no common structural theme behind these interactions ^29^. Because KMN complexes form and are stabilized in mitosis ^19,40^, it is important to assess if cooperative allosteric interactions favor the establishment of complete KMN assemblies, and to determine the precise path of force transmission within the KMN “antennas”. Phosphorylation of Mis12C by Aurora B also promotes binding to CENP-C and CENP-T, but the mechanistic basis of this regulation remains poorly understood ^22,23^. Finally, the structure of ZWINT is unknown, and so is the mechanism of the tight interaction with KNL1 that allows ZWINT to become incorporated into the KMN network.

Here, we shed light on these questions by reporting a cryo-EM reconstruction of a KMN subcomplex incorporating all interaction interfaces relevant to KMN assembly, which we validate with a combination of biophysical and cell biological experiments. Using model fitting and AlphaFold2 modelling, we generate a molecular model of the entire KMN network and discuss its properties and implications for kinetochore assembly and function.

## Results and Discussion

### Structure determination of the human KMN network

We reconstituted an 8-subunit KMN network comprising the full length Mis12 complex (Mis12C; Figure 1a), a dimer encompassing residues 109-197 of SPC24 (SPC24^109-197^) and residues 92-224 of SPC25 (SPC25^92-224^), ZWINT, and the C-terminal region of KNL1 (residues 2021-2342; note that numbering refers to isoform 1 of the *CASC5* gene that encodes KNL1). Elution from a size-exclusion chromatography column indicated a stoichiometric and monodisperse complex (Figure 1b).

The resulting KMN sample was processed for cryo-EM data collection, and a reconstruction was calculated to a resolution of 4.5 Å as described in Figure 1 – Supplement 1a-d. The map (Figure 1c) includes density for the entire Mis12C, the globular RWD domains of the SPC24/SPC25 dimer, and the tandem RWD domains (RWD^N^ and RWD^C^, Figure 2a) of KNL1, thus encompassing all interfaces that are relevant for the assembly of the KMN network. The region of KNL1 and ZWINT preceding the RWD domains are expected to interact in a heterologous helical coiled-coil (see below), but are not clearly distinguishable in our maps. We generated a molecular model of the KMN network by fitting previously determined high-resolution structures of the individual modules as discussed in Methods, complemented where necessary with AlphaFold2 (AF2) predictions, and refining this model against the experimental Cryo-EM map.

**Figure 2.**
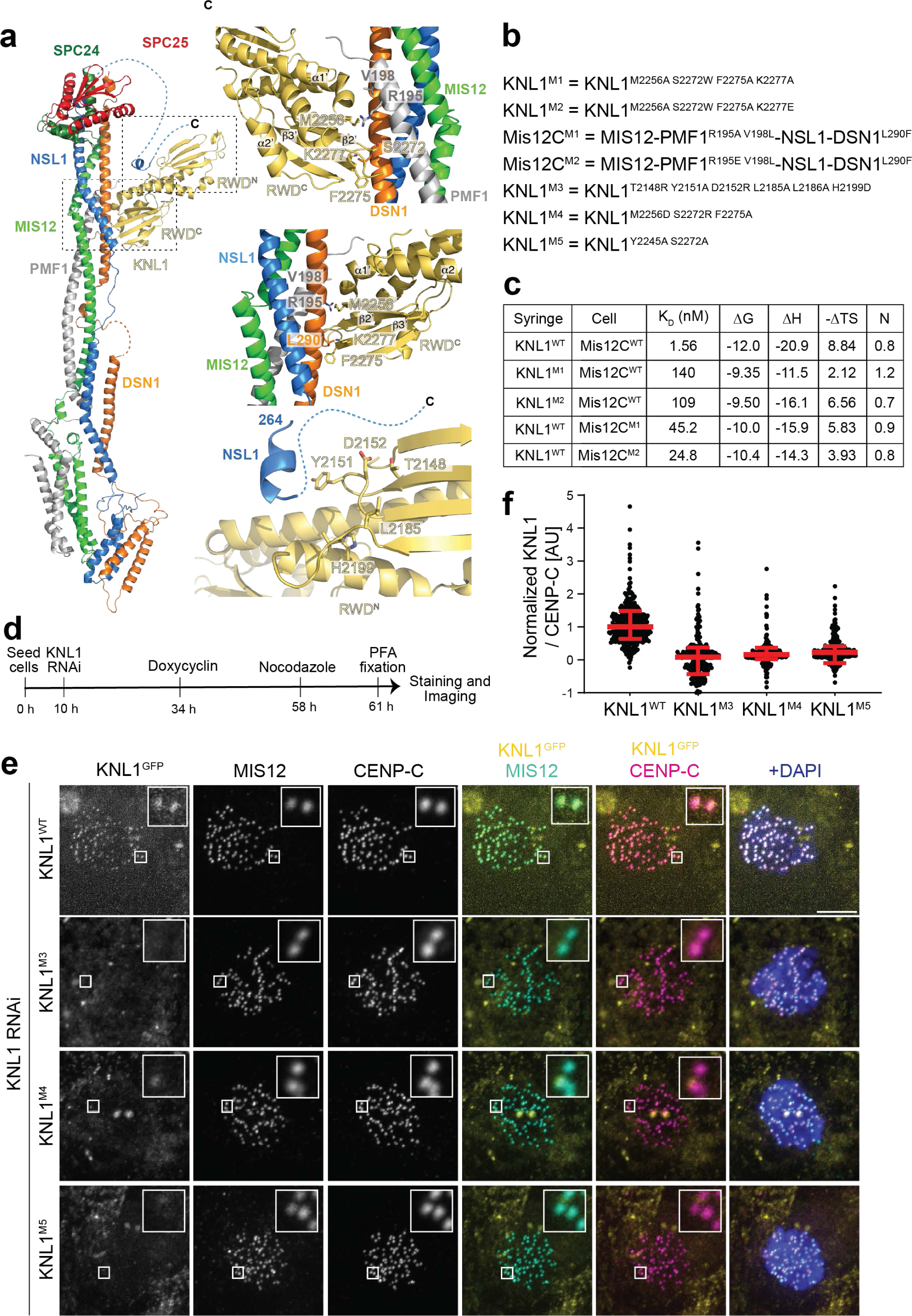
The Mis12C/KNL1 interaction. (**a**) Cartoon model of the minimal KMN network and closeups describing the Mis12C/KNL1 interface. (**b**) List of seven mutants chosen for biochemical or biological validation. (**c**) Table summarizing the results of the indicated ITC titration experiments. (**d**) Schematic representation of protocol for cell biology experiment shown in panel **e**. (**e**) Representative images of cells depleted of endogenous KNL1 and expressing the indicated EGFP-KNL1^2026-2342^ construct. Scale bar = 5 µm. (**f**) Quantification of the experiment in panel **e**. Red bars represent median and interquartile range of normalized kinetochore intensity values for KNL1^WT^ (n = 1037), KNL1^M3^ (n = 586), KNL1^M4^ (n = 760), KNL1^M5^ (n =566) from three independent experiments.

### Overall appearance of KMN network

The Mis12c is elongated, with a long axis of approximately 22 nm (Figure 1d). It starts with two globular heads, head1 and head2, each a four-helix bundle. Head1 and head2 respectively encompass the N-terminal α1-α2 helices of PMF1/MIS12 and DSN1/NSL1. The Mis12C then continues with a compact parallel helical bundle of the four subunits, the stalk. The SPC24/SPC25 dimer, consisting of two tightly interacting RWD domains, caps the complex near the C-terminal tails of the Mis12C stalk, engaging in tight interactions with the C-terminal regions of DSN1 and NSL1. The RWD^C^ of KNL1 docks against the lateral surface of the stalk, approximately 5 nm from the upper end, engaging the DSN1 and PMF1 subunits. It also captures a C-terminal motif of NSL1 that is connected to the stalk through a flexible (and largely invisible) linker.

In the KMN structure, the two Mis12C heads are closely packed against each other (Figure 1 – Supplement 2a). Comparison with a previously determined crystal structure of the complex of the Mis12C with CENP-C indicates that the closed conformation of the heads we observe is incompatible with CENP-C binding. In complex with CENP-C, head2 sways out of position, a conformational change that resolves a predicted steric clash with CENP-C (Figure 1 – Supplement 2a). Indeed, by binding at the interface of head1 and head2, CENP-C also pushes head1 and the stalk away from each other, so that the closed conformation of the unbound complex is also comparatively more compact, as shown by a superposition (Figure 1 – Supplement 2a). Aurora B kinase phosphorylates Ser100 and Ser109 (and possibly Ser95) on DSN1 to facilitate binding of CENP-C and CENP-T to Mis12C ^19-24^. Deletion of a loop region holding these residues bypasses this requirement, suggesting that the ground state of the loop stabilizes the closed conformation of the heads, and that phosphorylation facilitates their release into an open conformation. The experimental map is not sufficiently well-resolved in this region to pinpoint the exact position of the DSN1 phosphorylation loop, but AF2 provides a robust prediction for this region (Figure 1 – Supplement 2b). Specifically, the strongly positively charged Dsn1^80-112^ segment is predicted to bind at a negatively charged interface of head1 and head2, opposite to the CENP-C binding site, “gluing” the two heads together (Figure 1 – Supplement 2b). Phosphorylation may therefore weaken the closed state of the heads by modifying the charge balance in this region.

### The Mis12C/KNL1 interaction

Mis12C engages residues 282-304 of DSN1 and residues 188-204 of PMF1 (near the C-terminal ends of these subunits) to interact with the KNL1 RWD^C^. The strongly kinked end of the α1’ helix (superscripts indicate secondary structure elements in RWD^C^ and lack thereof in RWD^N^, see Figure 2 – Supplement 1a) and the β2’-β3’ loop of RWD^C^ insert into a guide generated by the intertwined PMF1 and DSN1 helices (Figure 2a). A second crucial contact, which mainly engages RWD^N^, captures the C-terminal segment of NSL1 (Figure 2a and Figure 2 - supplement 1b). This part of the structure is closely related to a previously determined crystal structure of the KNL1 C-terminal region with an NSL1 peptide (PDB ID 4NF9; ^29^). Besides the interaction with the KNL1 C-terminal region, AF2 modelling also suggests that the NSL1 tail is further stabilized by the terminal part of the ZWINT/KNL1 coiled-coil, as explained more thoroughly below. Thus, RWD^N^ and RWD^C^ respectively specialize in the interaction with the C-terminal region of NSL1 and with the DSN1 and PMF1 helices in the stalk.

Collectively, the bipartite interaction buries over 2000 Å^2^ of protein surface. The extensive interface showcases a combination of hydrophobic, hydrophilic, and charge-charge interactions. Using isothermal titration calorimetry (ITC), we determined the dissociation constant (K_D_) of the Mis12C/KNL1^2026-2342^ interaction to be ∼1.6 nM (Figure 2c and Figure 2 – Supplement 1c), in line with previous measurements ^22,28^. We repeated binding measurements with selected mutants predicted to affect the Mis12C/KNL1 interface (Figure 2b-c and Figure 2 – Supplement 1d-g). All mutants disrupted the interaction, causing a severe increase of the K_D_ (from ∼12- to ∼100-fold; Figure 2c).

In the hierarchy of kinetochore assembly, Mis12C is a platform for KNL1 recruitment. To test the effects of mutations on *in vivo* localization of KNL1, we applied the pipeline schematized in Figure 2d to deplete endogenous KNL1 by RNAi (Figure 2 – Supplement 1h) and replace it with stable transgenes expressing KNL1^2026-2342^ fused to EGFP to monitor kinetochore localization. The wild type KNL1 fragment localized normally to kinetochores, but three mutants (KNL1^M3^, KNL1^M4^, and KNL1^M5^; Figure 2c) designed to impair binding to Mis12C failed to demonstrate any level of kinetochore localization. Conversely, the levels of Mis12C were unaffected (Figure 2e-f), indicating that the mutated residues on KNL1 are important for binding to Mis12C at kinetochores.

### The Mis12C/Ndc80C interaction

The C-terminal regions of DSN1 and NSL1 mediate interactions with the SPC24/SPC25 dimer required to recruit the Ndc80C onto the kinetochore^28^. As they emerge from the end of the stalk with their α4 helices, both the DSN1 and NSL1 chains make a sharp turn to interact with SPC24/SPC25 (Figure 3a). Also in this case, the interaction is extensive, collectively burying ∼2500 Å^2^. The interaction of DSN1 with SPC24/SPC25 is closely reminiscent of the interaction of CENP-T with SPC24/SPC25 (^15^; PDB ID 3VZA). Specifically, the position and binding mode of the DSN1 α5 helix is very similar to that of an equivalent helix of CENP-T in complex with SPC24/SPC25 (Figure 3 – Supplement 1a). Indeed, the two helices in DSN1 and CENP-T have related sequences that alternate hydrophobic and positively charged residues ^22^.

**Figure 3.**
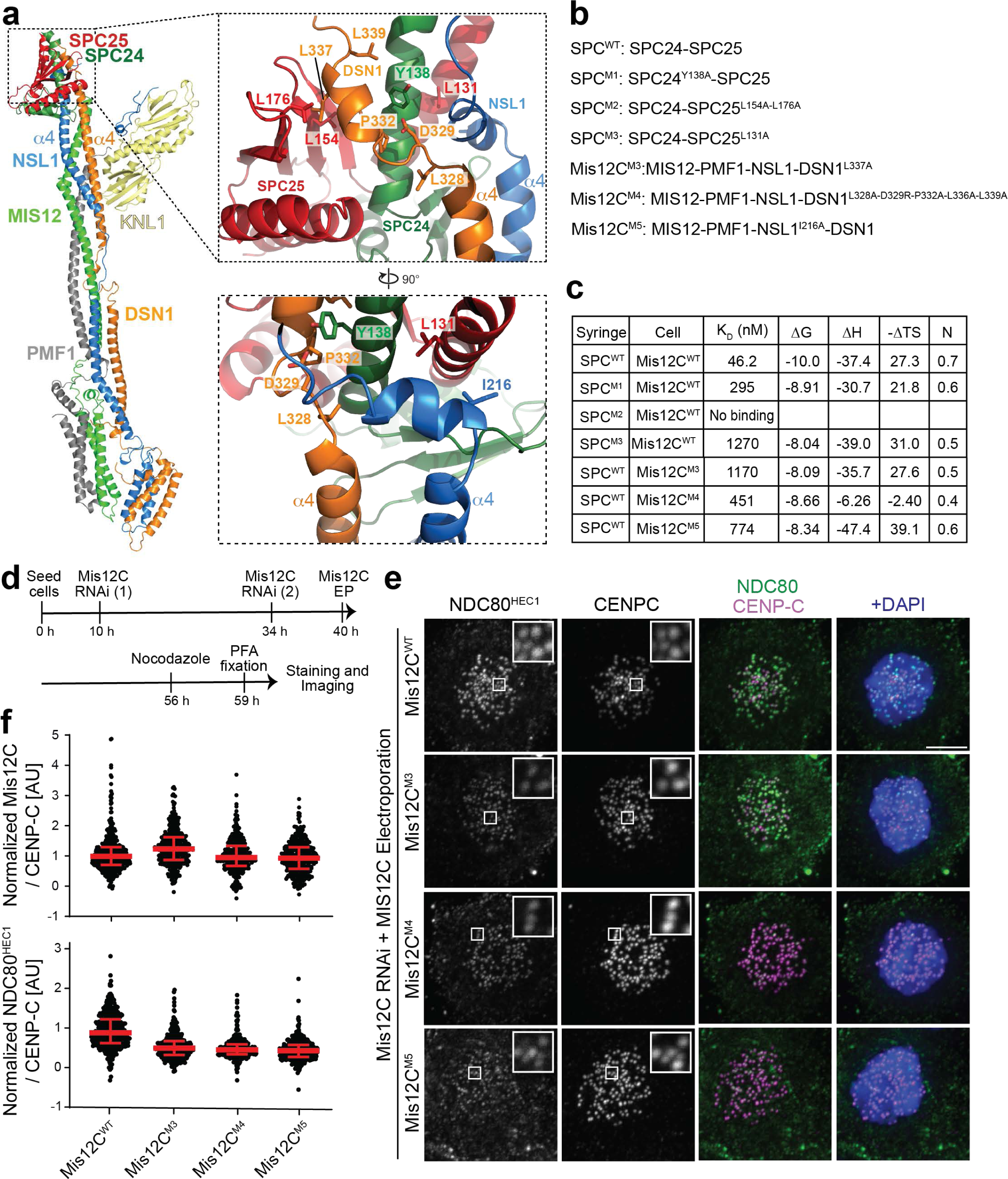
The Mis12C/Ndc80C interaction. (**a**) Cartoon model of the minimal KMN network and closeups describing the Mis12C/Ndc80C interface. (**b**) List of seven mutants chosen for biochemical or biological validation. (**c**) Table summarizing the results of the indicated ITC titration experiments. (**d**) Schematic representation of protocol for cell biology experiment shown in panel **e**. (**e**) Representative images of cells depleted of endogenous Mis12C and electroporated with the indicated Mis12C constructs. The levels of Ndc80C were monitored. Scale bar = 5 µm. (**f**) Quantification of the experiment in (**e**). Red bars represent median and interquartile range of normalized kinetochore intensity values for Mis12^WT^ (n = 365), Mis12^M3^ (n = 355), Mis12^M4^ (n = 298), Mis12^M5^ (n = 358) from three independent experiments.

Measured by ITC, the K_D_ of the interaction of SPC24/SPC25 with Mis12C was ∼46 nM (Figure 3c). Also using ITC, we previously measured a smaller K_D_ (4 nM) for the interaction with the entire Ndc80C ^28^, suggesting a slight decrease of binding affinity when using SPC24/SPC25. Two mutant constructs affecting residues at the interface with DSN1, SPC24^Y138A^/SPC25 and SPC24/SPC25^L154A-L176A^ (indicated as SPC^M1^ and SPC^M2^, respectively) had strongly reduced or non-measurable binding affinity for the wild-type Mis12C (Figure 3b-c). Similarly, mutants affecting hydrophobic residues in the DSN1 α5 helix, including Mis12C^M3^ and Mis12C^M4^ (Figure 3b-c), reduced the binding affinity for the SPC24/SPC25 complex by factors of ∼20 and ∼10, respectively. When reintroduced in HeLa cells depleted of endogenous Mis12C (following the protocol schematized in Figure 3d) the same mutants recruited Ndc80C sub-optimally, causing a ∼50 % reduction of the kinetochore levels of endogenous Ndc80C (Figure 3e-f).

NSL1 interacts with SPC24/SPC25 by engaging the PVIHL motif (residues 209-213) at the end of its α4 helix, previously shown to be essential for the interaction with the Ndc80C ^28^. A short α5 helix completes the interaction (Figure 3a and Figure 3 – Supplement 1a). Mutations of hydrophobic residues at the interface of the NSL1 α5 helix and SPC25 (listed as SPC^M3^ and Mis12C^M5^) reduced the binding affinity of Mis12C and SPC24/SPC25; the Mis12C^M5^ mutant had a similar disruptive effect on Ndc80C kinetochore recruitment as that described above for the Mis12C^M3^ and Mis12C^M4^ mutants. Thus, collectively, the mutational analysis was consistent with the interactions of the Mis12C with KNL1 and with the SPC24/SPC25 moiety of the Ndc80C revealed by our structural work.

### Mutual influence of KNL1 and Ndc80C on their kinetochore localization stability

The binding sites on Mis12C for KNL1 and for Ndc80C are clearly demarcated, and KNL1 and Ndc80C do not appear to interact directly. In previous work, however, we observed that individual depletions of KNL1 or Ndc80C had reciprocal effects on kinetochore localization ^41^. Indeed, depletion of KNL1 negatively affected the levels of kinetochore NDC80 (found to settle on approximately 80 % of control levels). Conversely, depletion of the Ndc80C caused an even more significant reduction in the levels of kinetochore KNL1 to 60 % of control levels (Figure 4a-b). Thus, KNL1 and the Ndc80C appear to influence each other’s stability at the kinetochore.

**Figure 4.**
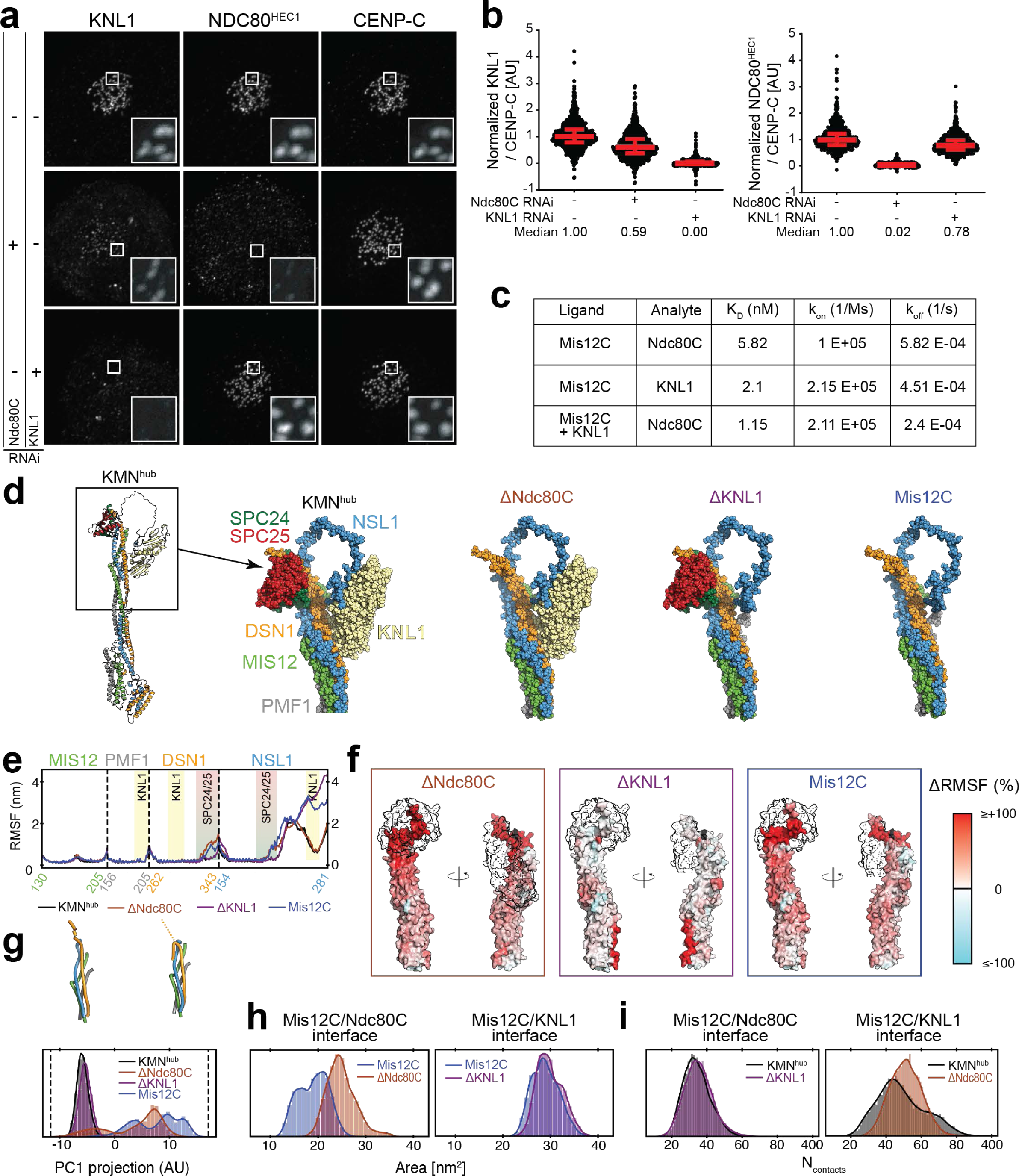
Dynamic binding interdependency in the Mis12-Knl1-Ndc80 complex. (**a**) Representative immunofluorescence analysis of the indicated kinetochore proteins upon RNAi-mediated depletion of Ndc80C subunits or KNL1 (see Methods). Scale bar = 5 µm. (**b**) Quantification of experiments displayed in **a**. Red bars represent average and interquartile range of normalized kinetochores intensity values for Ndc80C RNAi + KNL1 RNAi (n = 783), Ndc80C RNAi (n = 907), KNL1 RNAi (n = 848) from three independent experiments. (**c**) Summary of BLI experiments shown in Figure 4 – Supplement 1a-c. (**d**) Minimal fragment (KMN^hub^, black box) containing key interacting partners considered for MD simulations. Mis12C was truncated approximately at the middle of the helical bundle. In addition, the SPC24/SPC25 and KNL1 RWD headgroups of Ndc80C and KNL1, respectively, were considered. Summary of the four systems simulated (shown as spheres). (**e**) Per-residue root mean square fluctuations (RMSFs) of the Mis12C segments under four different conditions (legend at the bottom). The four helical elements are separated by the dashed lines and their sequence coverage is shown on the X-axis. The colored rectangles indicate the binding site of the SPC24/25 domains (red/green) and KNL1 (yellow). (**f**) Change in RMSF of the systems lacking at least one of the interaction partners relative to the system with both interaction partners bound (ΔRMSF=[RMSF(X)/RMSF(KMN^hub^)]^-1^, with X indicated at the top of each panel). The change is mapped on the surface of the Mis12C fragment according to the color scale (right). Residues that displayed an increased (reduced) RMSF in the absence of a binding partner are colored in red (cyan). The black outlines indicate the binding positions of the SPC24/SPC25 (*left*) and KNL1 domains (*right*). (**g**) Principal component analysis retrieved a main principal component (PC1) related to the bending motion of the C-terminus of the DSN1 domain (see cartoon representations). This component explained 33% of the positional fluctuations of the rigid part of the studied Mis12C fragment. Projection of the MD trajectories onto this PC component indicates whether the DSN1 unit adopted either a straight or a bent conformation. The normalized distribution of such projection is shown for the four studied systems (arbitrary units and same color as in **d**-**f**). (**h**) Normalized distribution of the exposed surface area of the Mis12C/Ndc80C (*left*) or Mis12C/KNL1 (*right*) binding interfaces for the indicated systems. (**i**) Normalized distribution of the number of atomic contacts (N_contacts_), between the Mis12C/Ndc80C (*left*) or Mis12C/KNL1 (*right*) binding interfaces is presented for the indicated systems.

To corroborate this conclusion with biochemical experiments, we used Bio-Layer Interferometry (BLI) to measure the binding affinity of the Mis12C for Ndc80C in presence or absence of KNL1^2026-2042^. Mis12C bound Ndc80C with a K_D_ of ∼6 nM (Figure 4c and Figure 4 – Supplement 1a-c), a dissociation constant almost 10-fold lower (indicative of tighter binding) than for the interaction of Mis12C with SPC24/SPC25 pair (Figure 3c). This result is consistent with the idea that the complete Ndc80C binds Mis12C with somewhat higher affinity than the SPC24/SPC25 subcomplex, as also in agreement with our previous measurements (as discussed above). However, we note that the two K_D_ measurements were performed with BLI and ITC, respectively, and therefore the differences in binding affinity may also reflect technical differences. We continued to use BLI to measure the strength of the MIS12/Ndc80C interaction in presence of KNL1^2026-2342^. The measured K_D_ of ∼1 nM (Figure 4c and Figure 4 – Supplement 1a-c) is consistent with the idea that KNL1 increases the binding affinity of Ndc80C for the Mis12C. The dissociation constant for the Mis12C/KNL1 interaction was ∼2 nM, almost identical to the value obtained by ITC (see Figure 2c). For technical reasons we could not address whether the addition of Ndc80C increased the binding affinity of the Mis12C/KNL1, an expectation raised by the biological experiments in Figure 4a-b.

### Molecular dynamics simulations reveal structural connectivity

To address the molecular underpinnings of interdependent binding of KNL1, Ndc80C and Mis12C, we conducted molecular dynamics (MD) simulations (see Methods). For computational efficiency, we considered a reduced fragment (“KMN^hub^”) containing a truncated Mis12C helical bundle and the SPC24/SPC25 and KNL1 binding headgroups (Figure 4d). In three additional sets of simulations, we removed SPC24/25 (ΔNdc80C), KNL1 (ΔKNL1), or both (Mis12C) (Figure 4d). Each system was simulated five times for 500 ns for a total cumulative sampling of 2.5 μs per system. Despite the truncation, the Mis12C helical bundle of KMN^hub^ was stable and SPC24/25 and KNL1 remained stably bound to it (Movie S1), with a root mean square deviation (RMSD) from the initial conformation levelling off at values smaller than 1 nm (Figure 4 – Supplement 1d). Omission of SPC24/SPC25, KNL1, or both increased the flexibility of the DSN1 and NSL1 segments without significantly impacting MIS12 and PMF1 (Figure 4e-f). Structural fluctuations were maximal in the regions implicated in Ndc80C and KNL1 binding, which were more rigid in association with the cognate ligands, and highly flexible upon their removal.

We inspected more closely the dependence between binding partners. (Un)binding of the SPC24/25 domains did not affect the RMSF of the KNL1 binding site and, reciprocally, (un)binding of KNL1 did not alter the RMSF of the SPC binding site (Figure 4e-f). Principal component analysis revealed that the DSN1 C-terminal region adopted a straighter orientation when the SPC24/SPC25 RWD domains were bound to it, irrespectively of KNL1 binding (see distributions along the 1^st^ principal component for KMN^hub^ and ΔKNL1 systems in Figure 4g). On the contrary, this segment of DSN1 adopted a broader range of conformations, including bent ones, when SPC24/SPC25 was not bound (ΔNdc80C and Mis12C in Figure 4g). Remarkably, in these systems where Ndc80C was omitted, KNL1 caused NSL1 to sample straight conformations reminiscent of those observed when NSL1 is bound to Ndc80C (shown by minor but significant peaks corresponding to straight orientations in the ΔNdc80C projection in Figure 4g). This tendency to stabilization was also evident near the Mis12C/Ndc80C interface, whose residues were significantly more solvent-exposed in presence of KNL1 (ΔNdc80C system) than in its absence (Mis12C system) (Figure 4h, *left*). Thus, our simulations suggest that binding of KNL1 shifts Mis12C into an unshielded straight conformation that prepares it for binding to SPC24/SPC25. The effect of Ndc80C on the KNL1-binding site in the simulations was less pronounced. The flexibility of the PMF1 and DSN1 helices at the KNL1 binding site remained largely unaffected by removal of any of the binding partners (Figure 4e-f and Figure 4g). Binding of SPC24/SPC25 did not alter the exposure of the KNL1 binding site (Figure 4h, *right*). Nevertheless, the number of atomic contacts at the Mis12C/KNL1 interface displayed a broader distribution in presence of SPC24/25 (Figure 4i, *right*), possibly indicating how SPC24/25 predisposes Mis12C to KNL1 binding.

Thus, collectively, the MD simulations provide a possible molecular explanation for the reciprocal dependence of KNL1 and Ndc80C on localisation and binding demonstrated in Figure 4a-c. When combined with our kinetic and thermodynamic characterization of the system (Figure 4c), this dependence is further captured by the thermodynamic cycle of Figure 4 – Supplement 1e, which depicts the binding of KNL1 or Ndc80C to Mis12C, in absence or presence of the reciprocal binding partner. The cycle implies that the Ndc80 complex thermodynamically stabilizes the Mis12C/KNL1 complex, and KNL1 reciprocally stabilizes the Mis12C/Ndc80C interaction.

### Limited plasticity of the Mis12C/Ndc80C connection

Thus, our studies suggest that the interactions of Knl1C and Ndc80C with Mis12C collectively stabilize the KMN. Whether these stabilizing effects are functionally related to microtubule-initiated force sensing within the KMN network is a question of great interest that will require analyses under controlled force application in an optical tweezer. Here, we asked if the connections of the Ndc80C with the Mis12C could be re-engineered on different Mis12C subunits. To this end, we generated several variants of the Mis12C in which the NSL1 and DSN1 C-terminal segments implicated in Ndc80C binding were grafted on the other two subunits, MIS12 and PMF1. All fourteen constructs we generated (listed in Figure 5 – Supplement 1a) co-eluted with KNL1^2026-2342^ in size-exclusion chromatography experiments (unpublished observations). On the other hand, only two constructs, Mis12C^18^ and Mis12C^19^ (Figure 5a) retained the ability to interact with Ndc80C (Figure 5b-c). In these constructs, grafting residues 227-281 of NSL1 (the NSL1 C-terminal segment downstream from the Ndc80C binding sites) onto either NNF1 or PMF1 was compatible with KNL1 binding. In no case we were able to swap the C-terminal segment of DSN1, or the C-terminal region of NSL1 comprising the Ndc80C binding region, onto another subunit of the Mis12C and retain binding to Ndc80C (Figure 5 – Supplement 1b).

**Figure 5.**
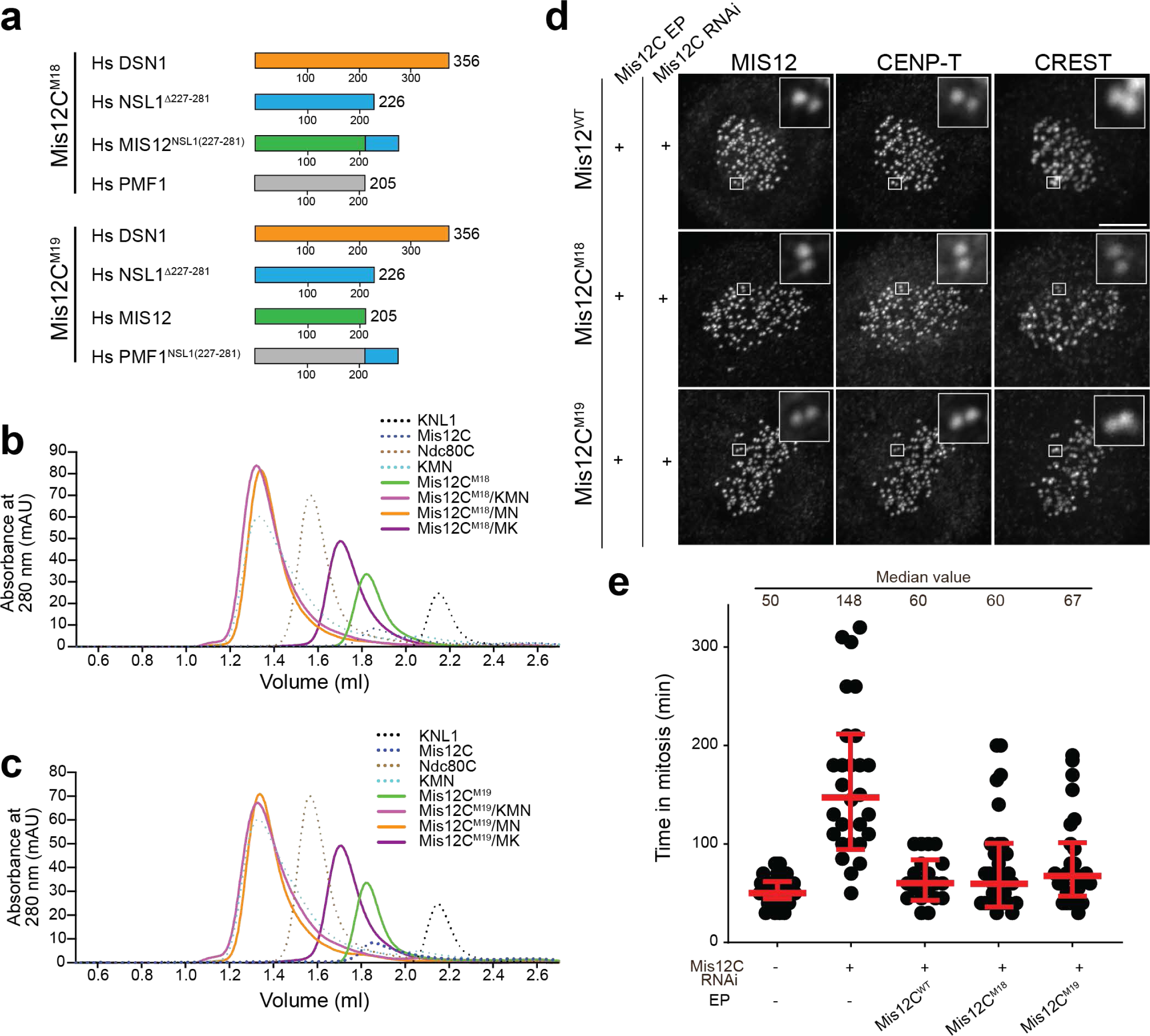
Swapping of Mis12C C-terminal tails. (**a**) Schematic representation of the Mis12C^M18^ and Mis12C^M19^ constructs. The detailed list of constructs is presented in Figure 5 – Supplement 1a. (**b**-**c**) Size-exclusion chromatography of the indicated biochemical species on a Superdex S200 5/150 column. (**d**) Representative images of cells depleted of endogenous Mis12C and electroporated with the indicated recombinant Mis12C species. Scale bar = 5 µm. (**e**) Time from nuclear envelope breakdown to chromosome decondensation in H2B-mCherry HeLa cells. Live cell movies were performed for 16 h. Each black dot represents a single cell. Median values of mitotic exit are reported for each condition.

When electroporated into HeLa cells depleted of endogenous Mis12C (ref. ^42^), Mis12C^WT^, Mis12C^M18^ and Mis12C^M19^ localized to kinetochores essentially indistinguishably (Figure 5d and Figure 5 – Supplement 1b) as expected on the basis of the experiments *in vitro* showing normal KMN assembly. Mis12C^WT^, Mis12C^M18^, and Mis12C^M19^ were all capable of restoring correct chromosome alignment and timing of mitotic exit, as revealed by their ability to suppress the mitotic arrest observed in cells depleted of endogenous Mis12C (Figure 5e). Thus, the segment of the NSL1 C-terminal tail downstream from the SPC24/SPC25 binding site can be grafted onto other Mis12C subunits without significant disruptions, an observation that suggests that communication between the binding sites for Ndc80C and KNL1 does not involve the NSL1 C-terminal region, but rather the helical stalk, as implied by our MD simulations.

### Structure and role of ZWINT

Density for ZWINT and for the KNL1 region preceding the RWD^N-C^ domains was not visible in our maps, even at low map contouring levels. We therefore opted to model the ZWINT/KNL1 complex using AF2 Multimer ^43^. The resulting model contains all ZWINT residues and residues 1880-2342 of KNL1 (KNL1^1880-2342^) (Figure 6 – Supplement 1) and shows high reliability scores (Figure 7 – Supplement 1a,e). The model consists of tightly intertwined ZWINT and KNL1 helices (Figure 6a). The model predicts ZWINT^76-220^ and KNL1^1960-2133^ to engage in a bent parallel coiled-coil that terminates into the KNL1 C-terminal globular domain. This coiled-coil is propped up by additional interactions with N-terminal fragments of both proteins, so that the coiled-coil’s initial segment is incorporated into a reinforced 4-helix bundle with short additional connecting helices. We used AF2 to carry out additional predictions of the Knl1C, including amphibian, avian, and yeast complexes (Figure 6 – Supplement

**Figure 6.**
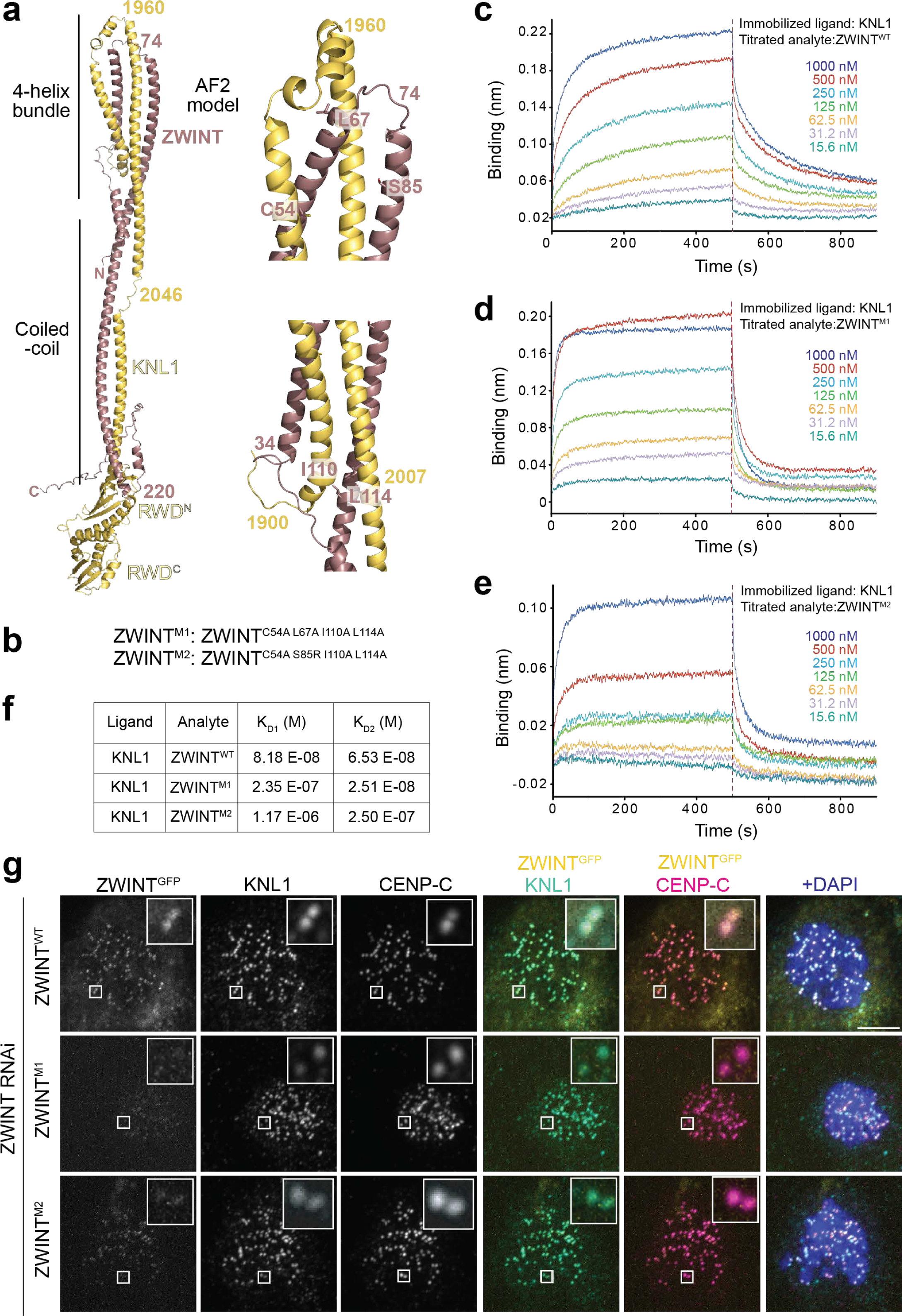
Model of the KNL1/ZWINT interaction and its validation. (**a**) AF2 model of the complex of KNL1^1880-2342^ and ZWINT with closeups showing position of complex-stabilizing residues. (**b**) Table describing two ZWINT mutants at interface residues. (**c**-(**e**) BLI traces for titration experiments with the indicated species. (**f**) Summary of binding titrations. (**g**) Representative images of HeLa cells depleted of endogenous ZWINT and expressing GFP fusions of the indicated ZWINT constructs from a stably integrated transgene. Quantifications of these experiments are displayed in Figure 6 – Supplement 1e. Scale bar = 5 µm.

**Figure 7.**
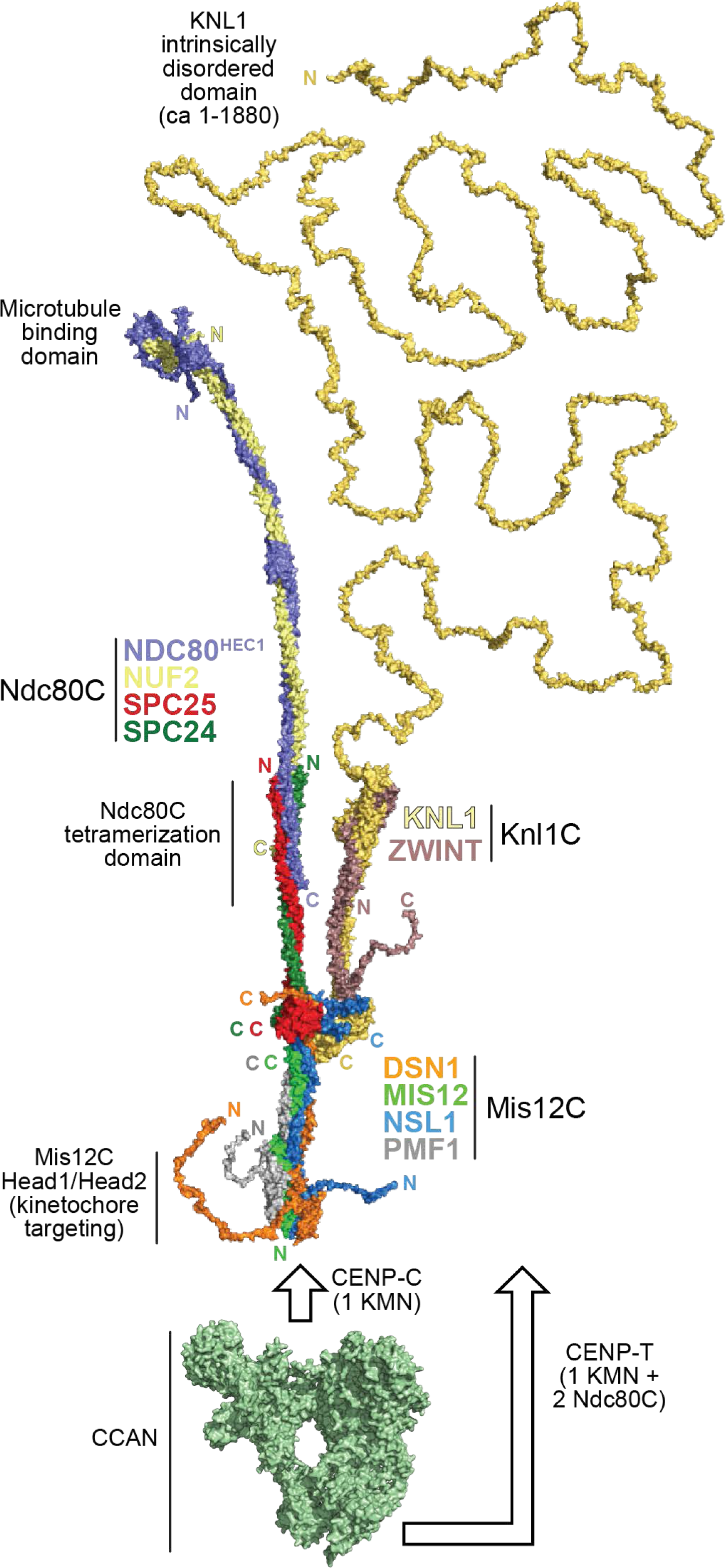
Model of the complete human KMN network. Surface representation of the human KMN hybrid model discussed in this paper (see Methods for description of approaches behind construction of the model and Figure 7 – Supplement 1 for AF2 quality scores). Each subunit is labelled with its color, and for each chain the position of N-and C-termini is indicated. The intrinsically disordered segment of KNL1 is included for reference. Shorter disordered tails are also present at the N-termini of NDC80^HEC1^, DSN1, and NSL1, as well as the C-termini of DSN1 and ZWINT. A surface representation of the 16-subunit CCAN complex (PDB ID 7QOO, displayed in green) ^80^ is also shown for reference and roughly in scale. In human, CCAN provides a point of attachment to the head1 of Mis12C through CENP-C. It recruits an entire KMN complex. CCAN also provides a second point of attachment to an entire KMN through CENP-T. Human CENP-T recruits two additional Ndc80C complexes through direct binding sites. See main text for details.

1a-d). Amphibian and avian Knl1C complexes are predicted to be closely related to the human complex. In both complexes, however, ZWINT is shown to have tandem RWD domains similar to those of KNL1, which appear to have been lost in humans. These observations strongly suggest that ZWINT and KNL1 are paralogs that have undergone significant functional and structural diversification after duplication from a common singleton. Yeast ZWINT (Kre28) is predicted to showcase a single RWD domain, and the overall appearance of its complex with KNL1 (Spc105) is much more compact relative to the other three modelled complexes (see also ^44^) (Figure 6 – Supplement 1d).

Based on the AF2 model of human Knl1C, we generated two ZWINT mutants (ZWINT^M1^ and ZWINT^M2^) harbouring point mutations predicted to destabilize the interface with KNL1 (Figure 6b). BLI measurements were best fitted by a 2-site binding model, and demonstrated that both mutants affected KNL1 binding (Figure 6c-f) When expressed from a stable transgene in cells depleted of endogenous ZWINT (Figure 2 – Supplement 1i), an ectopically expressed fusion of ZWINT^WT^ with EGFP localized robustly to kinetochores, whereas localization of both mutant constructs was very significantly impaired (Figure 6g; quantified in Figure 6–Supplement 1e). Conversely, the levels of endogenous KNL1 did not appear to be affected, suggesting that KNL1 can reach kinetochores independently of ZWINT. Collectively, these results validate the AF2 model and indicate that robust binding to KNL1 is necessary for ZWINT localization.

### A comprehensive model of the KMN network

Starting from the minimal core structure of the KMN network, we generated a comprehensive model of the KMN network by incorporation of ZWINT, of the entire KNL1 subunit, and of all Ndc80C subunits ^45^, with an overall molecular mass of 593 kDa (Figure 7). KNL1 is predicted to be intrinsically disordered essentially from its N-terminus until the point of entry into the helical domain (at residue 1880). The contour length of the disordered segment is predicted to be ∼700 nm (reference ^46^), several times longer than the predicted length of Ndc80C of ∼65 nm. A striking feature of the model is the prediction that the helical moiety of the Knl1C branch and the SPC24/SPC25 coiled-coil emerge from the Mis12C essentially in parallel. Despite absence of corroborating experimental information in our study, this conclusion seems reliable, as AF2 predictions of amphibian and avian Knl1C converge on similar models, where the angle of departure of the Knl1C coiled-coil from the RWD domains is largely fixed (although predictions of the avian complex also show bent conformations) (Figure 6 – Supplement 1b-c). This conclusion is also supported by experimental evidence in an independent accompanying study of the human outer kinetochore ^47^. Our structure may have failed to capture this feature due to the use in our reconstructions of a KNL1 fragment (Knl1^2021-2342^) that binds ZWINT robustly but may fail to fully stabilize the KNL1/ZWINT complex (refer to Figure 6a). The KMN network model suggests that the ends of the SPC24/SPC25 dimer and the beginning of the folded domain of the Knl1C may, with some likelihood, face each other at a relatively short distance. The functional implications of this configuration will have to be carefully analysed in future work. We speculate it may be related to spindle assembly checkpoint signalling, give the importance of Ndc80C and KNL1 in the recruitment of proteins involved in this pathway.

As already briefly mentioned, AF2 modelling predicts that the terminal segment of the ZWINT/KNL1 coiled-coil may contribute to the stabilization of the NSL1 C-terminal tail poised to interact with the KNL1 RWD^N^ domain (Figure 6 – Supplement 1f). Thr242 in this region of NSL1 has been recently shown to be a phosphorylation target of cyclin-dependent kinase that might destabilize the interaction of Knl1C with the kinetochore ^48^.

## Conclusions

We report the first comprehensive model of the human KMN network, the microtubule-binding apparatus of the kinetochore and a critical platform for mitotic checkpoint and error correction activities. Our conclusions substantially agree with, and are complementary to, those of an independent co-submitted study by Yatskevic, Barford and co-workers ^47^. The model sheds light on several fundamental aspects of KMN structure and function. First, it provides a glimpse of the two globular domains formed by the N-terminal region of the Mis12C, suggesting a mechanism for stabilization by DSN1 and switching to an open conformation through phosphorylation, thereby facilitating CENP-C binding. Second, the model reveals the molecular basis of the interactions of Knl1C and Ndc80C with Mis12C, which we further extensively validated experimentally. Third, the model indicates that ZWINT and KNL1 are paralogs. Their intertwined helices form an elongated domain that emerges in parallel to that of SPC24/SPC25. The helical domain of Knl1C may contribute to the stabilization of the interaction with NSL1 and may provide additional points of contact for checkpoint proteins known to require Ndc80C for kinetochore localization. The AF2 prediction indicates that the helical domain of KNL1 begins around residue 1880, and therefore earlier than the first residue of the KNL1 construct included in our experimental EM reconstruction. The KNL1 fragment we used for experimental structure determination is thermodynamically stable, mono-disperse, and interacts robustly with ZWINT. However, the model predicts that the KNL1 depletion might prevent assembly of the four-helix bundle (Figure 6a), possibly rendering this region of the structure more flexible and disordered. Fourth, molecular dynamics simulations and experimental perturbations indicate cooperative interactions of Ndc80C and Knl1C with Mis12C that stabilize the KMN assembly. Our simulations characterised the dynamics of the region where the Mis12C, Ndc80C, and Knl1C intersect, suggesting that Mis12C has flexible binding sites that become stabilised upon association of Ndc80C (via SPC24/25) and Knl1C (via KNL1). The increased binding affinity of the Mis12C/KNL1 complex for Ndc80C appeared to be determined by a slight increase in the association rate constant k_on_ relative to the Mis12C/Ndc80C interaction (Figure 4c). This, in turn, may reflect a more suitable-straighter and less shielded - binding conformation of the Mis12C, likely as a result of KNL1 binding. Increased stability of the product (i.e. the complex) reflected itself in a lower dissociation constant in presence of KNL1 (Figure 4c). The extremely low observed k_off_ rates (of the order of 10^-4^/s) are unreachable in the simulated time scale of microseconds and could not be resolved in our MD simulations. Nevertheless, the Mis12C/KNL1 complex interface may be more conformationally diverse and entropically more favourable when the Ndc80 protein domains were bound, as suggested by the more variable number of atomic contacts at the interface (Figure 4i).

In humans, CENP-C and CENP-T have each the ability to recruit one KMN complex (through interactions with Mis12C) to the kinetochore. Furthermore, the N-terminal region of human CENP-T provides binding sites for two additional isolated Ndc80C complexes ^13^. Thus, each CCAN complex is collectively predicted to recruit two KMN (each carrying Ndc80C and Knl1C) and two additional Ndc80Cs (i.e. four Ndc80C per CCAN). Microtubule binding appears to promote the assembly of Ndc80Cs into small clusters with enhanced potential to capture forces generated by depolymerizing microtubules ^45,49^. Our model of the KMN network provides a framework to understand the high-order organization of the kinetochore in future studies.

## Methods

### Cloning, expression, and purification of KNL1, MIS12, SPC24/SPC25, and ZWINT

All mutants in this study were generated by PCR-based site-directed mutagenesis. KNL1 constructs were expressed in *E. coli* BL21(DE3)-Codon-plus-RIPL in Terrific Broth supplemented with Chloramphenicol, Ampicillin, and 0.2 mM IPTG at 18°C for 16 h. Cells were pelleted down, washed once in PBS, and stored at −80°C for future use. All purifications were carried out on ice or at 4°C unless stated. For purification, frozen cells were removed from −80°C storage, thawed, and resuspended in GST lysis buffer [50 mM HEPES, pH 8.0, 150 mM NaCl, 2 mM TCEP, 10% Glycerol, 0.5 mM PMSF, DNase I, Protease-inhibitor mix HP plus (Serva)]. Cells were lysed by sonication and centrifuged at 108,000x*g* for 1 h. The cleared lysate was mixed with 5 ml Glutathione-agarose beads (5 ml of resin for 2 l of expression) and incubated on a roller overnight. The next day, beads were washed with 30 column volumes of wash buffer (lysis buffer without DNase and protease inhibitors). Beads were resuspended in wash buffer containing TEV protease and incubated overnight. Flow-through was passed through the Ni-beads to trap TEV protease and was dialyzed in GST SEC buffer [50 mM CHES, pH 9.0, 50 mM NaCl, 2 mM TCEP, 5% Glycerol]. The dialyzed protein was concentrated and injected into a HiLoad S75 16/600 column (GE Healthcare) pre-equilibrated with GST SEC buffer. Elution fractions were analyzed through Tricine-SDS-PAGE, and relevant fractions were pooled, concentrated, flash-frozen in liquid nitrogen, and stored at −80°C. All mutants of ZWINT were purified using the same buffers, conditions, and columns of KNL1 purification except 3C protease was added to remove the GST tag instead of TEV protease. The pBIG1A vector of the Mis12C was generated by combining expression cassettes of MIS12, PMF1, NSL1, and DSN1^SORT-HIS^. Bacmid from this vector were transfected in Sf9 insect cells to generate baculoviruses. Protein expression was carried out in Tnao38 insect cells. Cells were harvested after 72 h of transfection and stored at −80°C. For protein purification, cells were resuspended in Talon lysis buffer [50 mM HEPES, pH 8.0, 400 mM NaCl, 20 mM Imidazole, pH 8.0, 2 mM TCEP, 10 % Glycerol, 0.5 mM PMSF, DNase I, Protease-inhibitor mix HP plus (Serva)], lysed by sonication and centrifuged at 100,800x*g* for 1 h to clear the lysate. The lysate was filtered and added to 15 ml of Talon beads pre-equilibrated with Talon lysis buffer and incubated overnight. The beads were washed with 15 column volumes of Talon wash buffer I (lysis buffer without DNase and protein inhibitors) and then 3 column volumes of Talon wash buffer II (50 mM HEPES, pH 8.0, 400 mM NaCl, 30 mM Imidazole, pH 8.0, 2 mM TCEP, 10 % Glycerol). 20 ml of Talon elution buffer (Talon wash buffer I + 300 mM Imidazole) were added to beads and incubated for 2 h. Flow-through was collected, concentrated, and injected into HiLoad 16/600 Superose 6 column (GE Healthcare) pre-equilibrated with Mis12C SEC buffer (50 mM HEPES pH 8.0, 250 mM NaCl, 2 mM TCEP, 5% Glycerol). Elution fractions were analyzed by Tricine-SDS-PAGE and relevant fractions were pooled together, concentrated, flash-frozen in liquid Nitrogen, and stored at −80°C. SPC24/SPC25 and its mutants were expressed in BL21(DE3)-codon-plus-RIPL cells by induction of 0.4 mM IPTG at 18°C for 16 h. Cells were harvested, washed once in PBS, and stored at −80°C. Frozen pellets were resuspended in SPC lysis buffer [50 mM HEPES pH 8.0, 500 mM NaCl, 2 mM TCEP, 10% Glycerol, 1 mM EDTA, 0.5 mM PMSF, protease-inhibitor mix HP plus (Serva)], lysed by sonication, and the lysate cleared by centrifugation. The cleared lysate was added to Glutathione-Agarose beads pre-equilibrated with SPC wash buffer (SPC lysis buffer without DNase I and protease inhibitor) and incubated overnight. Beads were extensively washed with wash buffer and were incubated in SPC wash buffer containing 3C protease to cleave off the GST. Flow-through was collected, concentrated, and loaded into HiLoad S75 16/600 column (GE Healthcare) pre-equilibrated in SPC SEC buffer [50 mM HEPES pH 8.0, 250 mM NaCl, 2 mM TCEP, 5% Glycerol]. Relevant elution fractions were pooled, concentrated, flash-frozen in liquid Nitrogen, and stored at −80°C.

### Biotin and fluorescent labeling of proteins

C-terminal peptide conjugations in all the proteins were carried out by using calcium-independent Sortase 7M enzyme. The synthetic peptide used for biotinylation was GGGG^Biotin^ (Thermo Fisher Scientific) and for FAM labeling was GGGGK^FAM^ (Genscript). Reactions were performed at 4°C overnight with Sortase:Protein:Peptide in approximately 1:10:100 ratio in SEC buffer of relevant proteins. Excess peptide and Sortase were removed from the labeled protein by running the whole mixture on a size-exclusion chromatography column.

### Cell culture, siRNA transfection, and immunoblotting

HeLa cells expressing mCherry-H2B and DLD1 Flp-In-TREx cell lines were cultured in DMEM media (PAN Biotech) supplemented with 10% FBS (Clontech) and 2 mM L-glutamine (PAN Biotech) (DMEM complete media) in a humidifier chamber maintained at 37°C in presence of 5% CO_2_. Cell lines were regularly tested for mycoplasma contamination. The following siRNAs were used in this study: siMis12C (Dharmacon siMIS12 5’-GACGUUGACUUUCUUUGAU-3’; Sigma siDSN1 5’-GUCUAUCAGUGUCGAUUUA-3’, siNSL1 5’-CAUGAGCUCUUUCUGUUUA-3’); siKNL1 (Invitrogen HSS183683 5’-CACCCAGTGTCATACAGCCAATATT-3’, HSS125942 5’-TCTACTGTGGTGGAGTTCTTGATAA-3’, HSS125943 5’-CCCTCTGGAGGAATGGTCTAATAAT-3’); siZWINT (Sigma, 5’-GCACGUAGAGGCCAUCAAA-3’). RNAiMAX (Invitrogen) was used as a transfection reagent and transfections of siRNAs were carried out according to the manufacturer’s guidelines. For immunoblotting, cells were thawed on ice and resuspended in blot lysis buffer [75 mM HEPES pH 7.5, 150 mM KCl, 1.5 mM EGTA, 1.5 mM MgCl_2,_ 10 % Glycerol, 0.075% NP-40 and protease inhibitor cocktail (Serva)]. Cells were lysed by vigorously pipetting and were centrifuged at 16,000 g for 30 min at 4°C to clear lysate. Lysates were run in NuPAGE 4-12% Bis-Tris gradient gels and then proteins were transferred to Nitrocellulose membrane for further analysis. The following antibodies were used in this study for immunoblotting: anti-KNL1 (rabbit polyclonal, clone id: SI0787, Ossolengo, 1:1000), anti-Vinculin (mouse monoclonal, clone hVIN-1, Sigma-Aldrich, 1:10,000).

### Mammalian Cell electroporation

All electroporation experiments of this study were performed by using a Neon Transfection Kit [Thermo Fischer Scientific]. For electroporation, cells were collected after trypsinizing them. Cells were then washed with PBS, counted, and split between 2 and 3 million. Cells were washed once in PBS and resuspended in 90 µl of proprietary buffer R. Mis12C complex and its mutants were taken at a concentration of 40 µM for electroporation. Proteins were tawed and centrifuged at 16,000 g at 4°C for 10 min. Proteins were diluted with Buffer R in 1:1 ratio. Cells and proteins were mixed together and loaded into a 100 µl Neon Pipette Tip. The Pipette Tip was mounted to the electroporation adaptor chamber and cells were electroporated at 1005 V with two consecutive pulses of 35 msec. Cells were washed once in PBS, trypsinized to remove excess cell surface-bound proteins, and resuspended in DMEM complete media. Cells were then plated in 6-well plates with coverslips inside them for immunofluorescence analysis or in 24-well plates (Ibidi) for live-cell imaging.

### Generation of stable mammalian cell lines

Flp-In-T-REx DLD-1 osTIR1 cell lines were used to generate stable cell lines. Sequences encoding KNL1^2026-2342^ and ZWINT wild-type and mutants were cloned into a pCDNA5/FRT/TO-EGFP-IRES plasmid and co-transfected with pOG44 (Invitrogen), encoding the Flp recombinase, into freshly seeded cells using X-tremeGENE (Roche) transfection reagent. To obtain positive clones, cells were selected in DMEM complete media supplemented with hygromycin B (250 µg/ml, Thermo Fisher Scientific) and blasticidin (4 µg/ml, Thermo Fisher Scientific) for two weeks. Positive clones were pooled, expanded, and expressions of transgenes were verified by immunofluorescence and immunoblot analyses. Gene expression was induced by addition of 0.3 µg/ml of doxycycline (Sigma-Aldrich) for 24 h.

### Cell treatment, microscopy, immunofluorescence detection, and live cell imaging

For immunofluorescence analysis, cells were treated with Nocodazole (3.3 µM) or Nocodazole and MG132 (10 µM) for 3 h before fixing them for staining. Cells were permeabilized with 0.5% Triton-X in PHEM buffer and fixed on coverslips with 4% paraformaldehyde. Cells were then blocked with 5% boiled donkey serum and stained with primary antibodies in a humidified chamber at room temperature for 2 h. Cells were washed with 0.1% Triton-X in PHEM buffer to remove excess primary antibodies and stained with secondary antibodies at room temperature for 1 h. DNA was stained with DAPI and coverslips were mounted on slides with Mowiol. The following primary and secondary antibodies were used in this study: NSL1 [mouse monoclonal, QL24-1 (25), in-house, 1:800], anti-CENP-C (guinea pig polyclonal, MBL-PD030, MBL, 1:1000), anti-HEC1 (human NDC80, mouse monoclonal, ab3613, Abcam, 1:3000), CENP-T (rabbit polyclonal, SI0822, in-house, 1:1000), KNL1 (rabbit polyclonal, SI0788, 1:500), goat anti-guinea pig Alexa Fluor 647 (Invitrogen A-21450), Goat anti-mouse Alexa Fluor 488 (Invitrogen A A11001), goat anti-mouse Rhodamine Red (Jackson Immuno Research 115-295003), donkey anti-rabbit Alexa Fluor 488 (Invitrogen A21206), donkey anti-rabbit Rhodamine Red (Jackson Immuno Research 711-295-152), goat anti-human Alexa Fluor 647 (Jackson Immuno Research 109-603-003). For immunofluorescence imaging, cells were imaged with a customized 3i Marianas system (Intelligent Imaging Innovations) equipped with an Axio Observer Z1 microscope chassis (Zeiss), a CSU-X1 confocal scanner unit (Yokogawa Electric Corporation), Plan-Apochromat 100x/1.4 NA objective (Zeiss), and an Orca Flash 4.0 sCMOS camera (Hamamatsu). Images were taken as a Z section, one channel at a time using Slidebook software. For live-cell imaging, cells were plated on 24-well imaging plates (Ibidi) and 1 h prior to starting movies, complete DMEM media was replaced with CO_2_ independent L-15 imaging media containing 10% FBS, with or without drugs. Images were acquired using a DeltaVision Elite System (GE Healthcare) equipped with an IX-71 inverted microscope (Olympus), a UPlanFLN 40x 1.3 NA objective (Olympus), and a pco.edge sCMOS camera (PCO-TECH Inc.) at 37°C. Live cell images were taken as a Z-scan starting from the bottom of the plate, one channel at a time per Z every 10 min intervals for 16 h. The softWoRx software was used to set up movies and further analysis. All images were visualized and processed using Fiji ^50^. Quantifications of intensities of protein signals were calculated using custom written script using the DAPI channel as a masking boundary. All protein signals were normalized to either CENP-C or CREST. To quantitate time in mitosis,16 h long movies were analyzed manually. Cells entering the mitosis were taken as entry points while cells exiting mitosis were taken as end points. The time differences between entry and exit points are represented as Time in mitosis in the analysis in Figure 5e.

### Isothermal Titration Calorimetry

All ITC experiments were carried out using PEAQ-ITC microcalorimeter (Malvern Preanalytical). All proteins were dialyzed overnight at 4°C in ITC buffer (50 mM HEPES, pH 8.0, 150 mM NaCl, 1 mM TCEP) to correct buffer mismatches. A typical ITC experiment consists of 19 injections of protein from the syringe, with the first injection of 0.4 µl followed by 18 injections of 2 µl with a spacing of 150 s to allow the heat change curve to hit again the baseline. Binding isotherms were integrated, corrected for offset, and the data were fitted in one-site binding equation in Microcal PEAQ-ITC analysis software.

### Biolayer Interferometry

All Biolyer Interferometry experiments were carried out on a FORTÉBIO BLI instrument. All proteins were dialyzed in BLI dialysis buffer (50 mM HEPES pH 8.0, 150 mM NaCl, 2 mM TCEP, 0.05 % Tween-20) to reduce buffer mismatch. Experiments were performed at 25°C by using Octet SA Biosensors. Biotinylated proteins (ligands) were loaded on sensors and different concentrations of analytes were used for binding to the bound ligand. Resulting curves were fitted either in 1:1 or 2:1 binding model to get association constants.

### KMN reconstitution

KNL1^2021-2342^, ZWINT, Mis12C, and SPC24^109-197^/SPC25^92-224^ were mixed together in equimolar concentration in KMN reconstitution buffer [50 mM HEPES pH 8.0, 150 mM NaCl, 2 mM TCEP pH 8.0] in ice for at least 1 h. The sample was injected in S200 5/150 column (GE Healthcare) pre-equilibrated with KMNZ reconstruction buffer and the peak fraction was taken for structural analysis.

### Analytical size-exclusion chromatography

Proteins were mixed in equimolar concentrations, incubated on ice for at least 2 h, centrifuged at 16,000 g for 20 min at 4°C and then injected into Superose 6 5/150 increase column (GE Healthcare) pre-equilibrated with analytical size-exclusion chromatography buffer (50 mM HEPES, pH 8.0, 150 mM NaCl, 2 mM TCEP). Typically, fractions of 80 µl were collected at a manufacturer’s recommended flow rate and analyzed by running in SDS-PAGE.

### Grid preparation and Cryo-EM data acquisition

Grids were prepared using a Vitrobot Mark IV (Thermo Fisher Scientific) at 13°C and 100% humidity. 4 μl of KMN complex at 0.3 mg/ml supplemented with 0.003% Triton were applied to glow-discharged UltrAuFoil R2/2 grids (Quantifoil). Excess liquid was removed by blotting (3.5 s at blot force −3), followed by vitrification in liquid ethane. Cryo-EM data were recorded on a Cs-corrected Titan Krios G2 microscope equipped with a field emission gun. 9,896 micrographs were acquired on a K3 camera (Gatan) in super-resolution mode at a nominal magnification of 105,000, corresponding to a physical pixel size of 0.68 Å/px and super-resolution pixel size of 0.34 Å/px, respectively (Figure 1 – Supplement 1a). A total of 63.9 e−/Å2 was distributed over 60 frames. The Bioquantum energy filter (Gatan) zero-loss slit width was set to 15 eV and the defocus range from −1.5 µm to −3.0 µm. Details on data acquisition and processing can be found in Table S1.

### Processing of Cryo-EM data

Data were pre-processed on the fly in TranSPHIRE ^51^, including motion correction and dose-weighting using MotionCorr2 ^52^, estimation of CTF parameters using CTFFIND4 in movie mode ^53^ and particle picking in crYOLO using a neuronal network trained on 20 representative micrographs of the dataset ^54^. 2,925,200 particles were picked by crYOLO, and 1,792,983 2-fold binned particles that had a crYOLO confidence value of 0.2 or higher were extracted in SPHIRE ^55^ with a box size of 256 pixels (Figure 1 – Supplement 1b). 2D classification was performed in CryoSPARC 4.2.1 ^56^ and 582,795 particles associated with high quality 2D classes were selected for 3D refinement in RELION 3.1.2 ^57,58^ using a reference volume created from another, independent dataset that was acquired using similar imaging settings, but with a volta phase plate to increase contrast. This initial 3D refinement reached 7.1 Å resolution. Particle polishing was performed in RELION and all further steps carried out in CryoSPARC. Polished particles were subjected to a second round of 2D classification. 430,553 particles assigned to high quality 2D classes were subjected to heterogeneous refinement using one good and three junk models. 230,597 particles classified into the class associated with the good model. Those were refined to a nominal resolution of 4.5 Å (Figure 1 – Supplement 1b-c), although the reconstruction does not extend to this resolution in all directions due to heavily preferred orientations that could not be alleviated by 2D class rebalancing or similar approaches (Figure 1 – Supplement 1d). DeepEMhancer ^59^ was used to create the final map that is displayed in the figures (Figure 1 – Supplement 1b).

### Molecular modelling

AlphaFold 2 (AF2) was used for all molecular modelling in the version AF2 Multimer 3.2.1 ^43,60^. Since AF2-multimer tends to artificially bend extended coiled coil models even more than the original AF2 ^61^, the models comprising the Ndc80 complex were predicted in segments. For the KMN model, three subcomplexes were predicted: model *i*) Ndc80c N-terminal region with NDC80^1-530^ and NUF2^1-368^; model *ii*) Ndc80c with NDC80^312-C^, NUF2^476-C^, SPC24, and SPC25; and model *iii*) Mis12c complex with ZWINT^FL^, KNL1^1880-C^ (isoform1 numbering) and SPC24/SPC25. These three models were stitched together using Ndc80^1-498^ and Nuf2^1-337^ of model *i*; NDC80^499-642^, NUF2^338-464^, SPC24^1-97^, and SPC25^1-91^ of model *ii*; and SPC24^98-197^, SPC25^92-224^ and all remaining chains from model *iii*. Thus, all relevant interfaces are predicted within a single AlphaFold model. The fragments were superimposed with PyMOL Molecular Graphics System, Version 2.5.4 Schrödinger, LLC.), and the stitching points within the alpha-helices were relaxed using alpha-helical geometry restraints within COOT ^62^. In the final model of the full length KMNZ (excluding the disordered segments), 90% of the residues had very reasonable pLDDT scores above 70; 77% had scores above 80, i.e. were predicted at least as “confident”; and 34% of the residues scored above 90, i.e. with very high confidence. Corroborating the high quality of the model, the “predicted alignment error” (PAE) plots consistently indicated well-defined relative positions of the domains to each other, including the tetramerization domain of Ndc80c that is composed of SPC24, SPC25, and the C-termini of NDC80 and NUF2. This model was used to create the Cryo-EM model by fitting into the experimental map, deleting the parts that are either not present in the constructs used for Cryo-EM experiments or invisible due to flexibility, and optimized iteratively by manual building on COOT ^62^ and real-space refinement in PHENIX ^63^. Panels showcasing structures were prepared using UCSF ChimeraX ^64^ or PyMOL. For the KNL1^1880-2342^/ZWINT complexes, complete models were predicted with AF2 Multimer 3.2.1. The sequence of chicken ZWINT, which was not available from the databases, was kindly provided by Tatsuo Fukagawa. For the human model (Accession IDs Q8NG31 (KNL1^1880-2342^) and O95229), 82% of residues have a pLDDT score over 70. In the other species the scores are 89% over 70 for *Xenopus laevis* (accession IDs A0A8J1LL48 (Knl1^2330-C^) and A0A8J1LF42), 80% over 70 for Gallus gallus (A0A8V0ZUL4 (Knl1^1500-C^)), and 65% over 70 for the *S. cerevisiae* model (SPC105^421-C^ P53148, Kre28 Q04431).

### Molecular dynamics (MD) simulations

The simulated system consisted of a truncated C-terminal part of the helical bundle of the Mis12C (residues 130–205 of MIS12, 156–205 of PMF1, 262–343 of DSN1, and 154–281 of NSL1) and the C-terminal headgroups of the SPC24 (residues 127-197), SPC25 (residues 126-224) and KNL1 domains (residues 2131-2342). Initial positions were taken from the cryo-EM structure determined in this study. The termini of the truncated fragments were capped with neutral groups. The protonation state of histidines was determined with PROPKA ^65^, through the APBS web server ^66^. The CHARMM36m force field ^67^ was used for the protein, the CHARMM-modified TIP3P model for the water molecules ^68^, and default CHARMM parameters for the ions. The Mis12C fragment was simulated with both the SPC24/25 and the KNL1 domains bound (system called KMN^hub^), without the SPC24/25 domains but with the KNL1 domain bound (ΔNdc80C), with the SPC24/25 domains but without the KNL1 domain bound (ΔKNL1), and alone (Mis12C). In all four cases, the protein complex was placed in a dodecahedron box, such that the distance between it and the periodic boundaries was at least 2 nm. The protein was surrounded by approximately 141000 water molecules. Sodium and Chloride ions were added at a concentration of 0.15 M and additional ions were added to neutralise the net charge of the protein. The solvated KMN^Hub^, ΔNdc80C, ΔKNL1, and Mis12C systems consisted of 434355, 434290, 434243, and 434071 atoms in total, respectively. The potential energy of the solvated systems was energy minimized by using the steep descent algorithm. Subsequently, the systems were thermalized, by maintaining both the volume and the temperature constant during 500 ps (NVT conditions). A solvent relaxation step of 1 ns followed, with both the pressure and temperature constant (NPT conditions). During these two equilibration steps, positions of the heavy atoms of the protein complex were harmonically restrained (elastic constant of the harmonic restraints: 1000 kJ/mol/nm^2^). After release of the position restraints, simulations of 500 ns were performed. Five independent simulation runs (after the energy minimization step) were carried out for each system for a cumulative simulation time in the production run phase of 2.5 μs per system. The first 100 ns of each production run was accounted as equilibration time and therefore discarded from the analysis.

The temperature was kept constant at a reference value of 310 K using Berendsen thermostat ^69^ during the NVT and NPT steps and the V-rescale thermostat ^70^ during production runs (with a coupling time constant of 1 ps). Pressure was maintained constant at 1 bar using the Berendsen ^69^ (NPT) and C-rescale ^71^ barostats (coupling time constant of 5 ps and reference compressibility of 4.5×10-5/bar). Both temperature and pressure were updated every 100 integration steps. Neighbours were treated according to the Verlet buffer scheme ^72^ with an update frequency of 20 steps and a buffer tolerance of 0.005 kJ mol^-1^ ps^-1^. Electrostatic interactions were calculated using the particle mesh Ewald algorithm ^73,74^, computing the Coulomb potential in the direct space for distances smaller than 1.2 nm and in the reciprocal space for larger distances. Short-range interactions were modelled by a Lennard Potential truncated at a distance of 1.2 nm. Bonds with hydrogen atoms in the protein complex were constrained via LINCS ^75^. In addition, both bond and angular degrees of freedom of water were constrained using Settle ^76^. Equations of motion were integrated using the Leap Frog algorithm at discrete time steps of 2 fs. Simulations were carried out using the GROMACS MD package ^77^ (2023 version).

Rigid-body translation and rotation of the centre of mass of the protein was removed prior to the simulation analyses, by least-squares superposition of the conformations at different simulation times with the initial conformation. The backbone atoms of the rigid part of the Mis12 complex, excluding the flexible loop of the NSL1 domain, and in the case of the RMSD calculation the SPC24/25 and KNL1 domains, were considered for the least-squares superposition. The following observables were extracted from the MD simulations. Unless stated otherwise, normalized histograms of the quantities of interest were calculated, after concatenating the 5 independent simulation replicas of each system. The root mean square deviation (RMSD) of the coordinates of the backbone atoms of all protein domains (except the highly flexible C-terminus of the NSL1 domain) was monitored, relative to the initial coordinates. The RMSD was computed using the GROMACS gmx rms tool. The root mean square fluctuation (RMSF) of each residue of the Mis12 complex was computed after concatenation of the 5 simulation runs of each respective system. The RMSF was computed with the GROMACS gmx rmsf utility. The relative change in ΔRMSF was defined as ΔRMSF=[RMSF(X)/RMSF(KMN^hub^)]-1, with X corresponding to the ΔNdc80C, ΔKNL1, and the Mis12C systems. Accordingly, ΔRMSF takes positive values if the RMSF of the X system (i.e. a system lacking at least one of the binding partners) is larger than that of the KMN^hub^ system. Reciprocally, ΔRMSF takes negative values if the RMSF of the X system is smaller. Principal component analysis, consisting on the calculation and diagonalization of the covariance matrix of the atomic positions ^78^, was carried out for the backbone atoms of rigid part of the Mis12C (i.e. discarding the flexible NSL1 C-terminus). To highlight the conformational changes due to the presence (or absence) of the binding partners, the MD trajectories of the four simulated systems were concatenated. The first principal component eigenvector explained ∼33% of the variation of the positions and was considered for further analysis. MD Trajectories were projected along this first principal component. This analysis was performed by means of the gmx covar and gmx anaeig tools. The solvent accessible surface area of the Ndc80 (i.e. the SPC24/25 domains) and the KNL1 (i.e. the Knl1 domain) binding sites on the Mis12C was computed. The area was calculated for the systems for which the binding partners were not present and therefore their binding sites were exposed, i.e. the ΔNdc80C, ΔKNL1, and Mis12C systems. The area was obtained by using the double cubi lattice method ^79^ probing, as solvent, a sphere of radius 0.14 nm. The GROMACS gmx sasa tool was used for this calculation. Mis12C residues at the binding interface were assumed to be those which were at a distance of 5 nm or less from the respective binding protein in the initial conformation of the KMN^hub^ system and they were identified with PyMOL (Schrödinger, L. & DeLano, W., 2020. PyMOL, Available at: http://www.pymol.org/pymol). Accordingly, the KNL1 binding site was composed of aminoacid segments 185–204 of PMF1, 283–305 of DSN1, and 255–273 of NSL1, and the SPC24/25 binding site of segments 313–342 of DSN1 and 197-224 of NSL1. The number of atomic contacts between heavy atoms of Mis12 and heavy atoms of the Ndc80C (SPC24/25) and Knl1C (KNL1) binding domains were obtained. Two heavy atoms were assumed to be in contact if they were at a distance equal or smaller than 0.35 nm. To avoid redundancy, contacts corresponding to the same atom in Mis12C were grouped and counted as one contact. The number of contacts was computed with the GROMACS gmx mindist option. Simulation analyses were carried out with our own in-house Bash and Python scripts. Protein conformations and movies were rendered with PyMOL.

### Data availability

Maps and coordinates will be deposited in the appropriate databanks before publication.

## Acknowledgements

We are grateful to Stefano Maffini and Nico Schmidt for help with microscopy experiments and data analysis, Oliver Hofnagel and Daniel Prumbaum for help in EM data collection, Sabrina Ghetti for critical reading of the manuscript, Tatsuo Fukagawa for sharing unpublished sequences, and Stanislau Yatskevich and David Barford for sharing unpublished results. A.M. gratefully acknowledges funding from the Max Planck Society, the European Research Council (ERC) Synergy Grant 951430 (BIOMECANET), the Marie-Curie Training Network DivIDE (project number 675737), the DGF’s Collaborative Research Centre 1430 “Molecular Mechanisms of Cell State Transitions”, and the CANTAR network under the Netzwerke-NRW program. MK, CA-S, and FG are grateful for financial support by the Klaus Tschira Foundation.

## Funding

Open access provided by the Max Planck Society

**Author contributions (following CRediT model) Conceptualization:** AM

**Data curation:** N/A

**Formal analysis:** TR, MK, CA, IRV

**Funding acquisition:** SR, FG, AM

**Investigation:** SP, TR, MK, CA, IRV

**Project Administration:** CA, FG, AM, SR, IRV

**Resources:** MT

**Supervision:** CA, FG, AM, SR, IRV **Validation:** CA, AM, SR, IRV **Visualization:** CA, AM, SP, TR, IRV **Writing – original draft:** AM

**Writing – review & editing:** CA, FG, AM, SP, TR, SR, IRV

### Ethics declarations

The authors declare no competing interests

## List of Supplemental Data

**Table S1**

**Figure 1 – Supplement 1**

**Figure 1 – Supplement 2**

**Figure 2 – Supplement 1**

**Figure 3 – Supplement 1**

**Figure 4 – Supplement 1**

**Figure 5 – Supplement 1**

**Figure 7 – Supplement 1**

**Movie S1**

**Figure 1 – Supplement 1.**
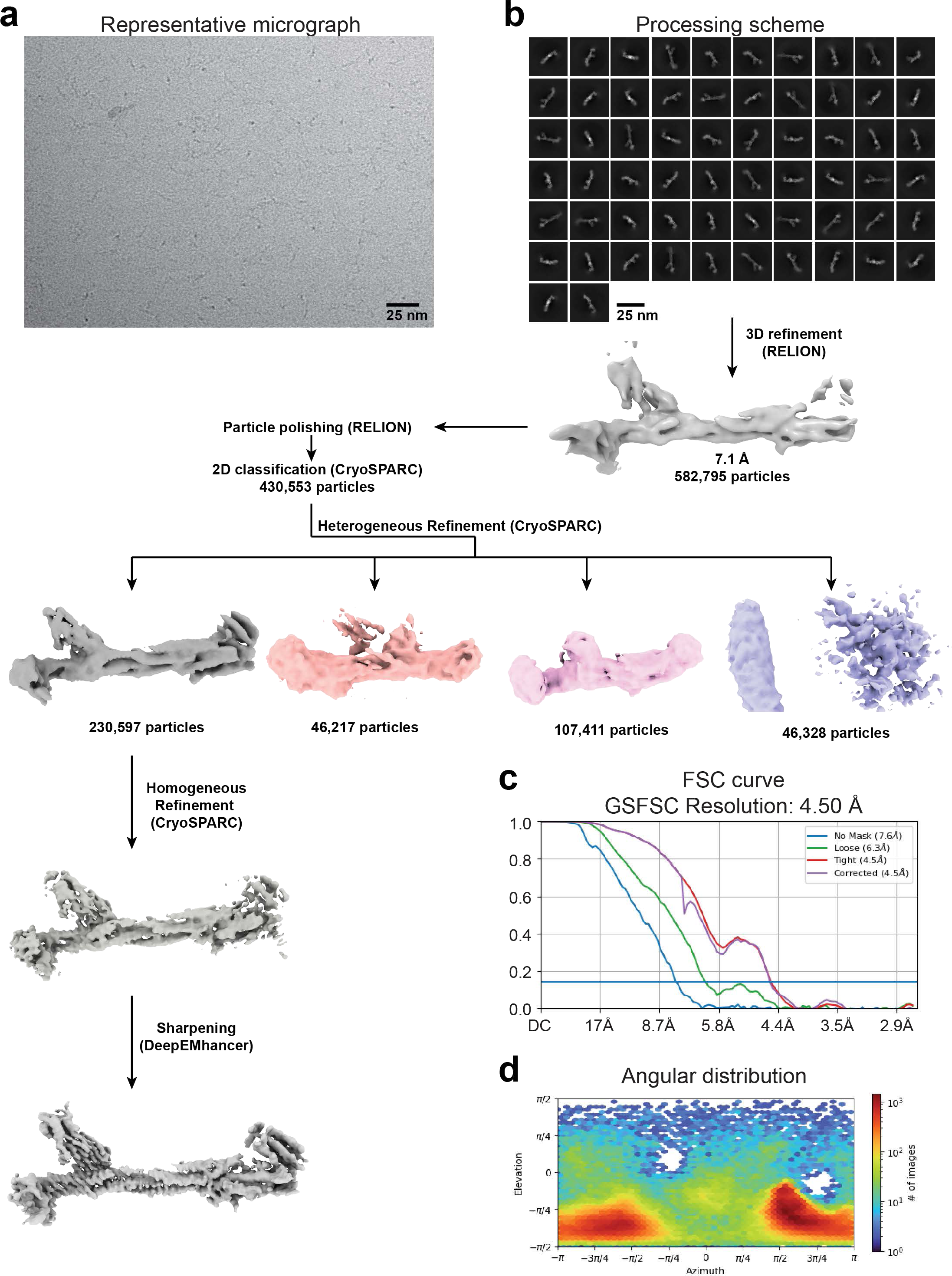
Cryo-EM data acquisition and processing strategy. (**a**) Representative micrograph of the KMN sample. (**b**) Outline of data processing procedure (see Methods). (**c**) Fourier shell correlation (FSC) plots between two independent half-maps without masking (blue) or masking with a loose (green) or tight (red) mask around the particle that have been automatically created by cryoSPARC. The purple curve corresponds to masking with the tight mask with correction by noise substitution. The horizontal blue line marks the 0.143 gold-standard FSC criterion. (**d**) Angular distribution of the particles.

**Figure 1 – Supplement 2.**
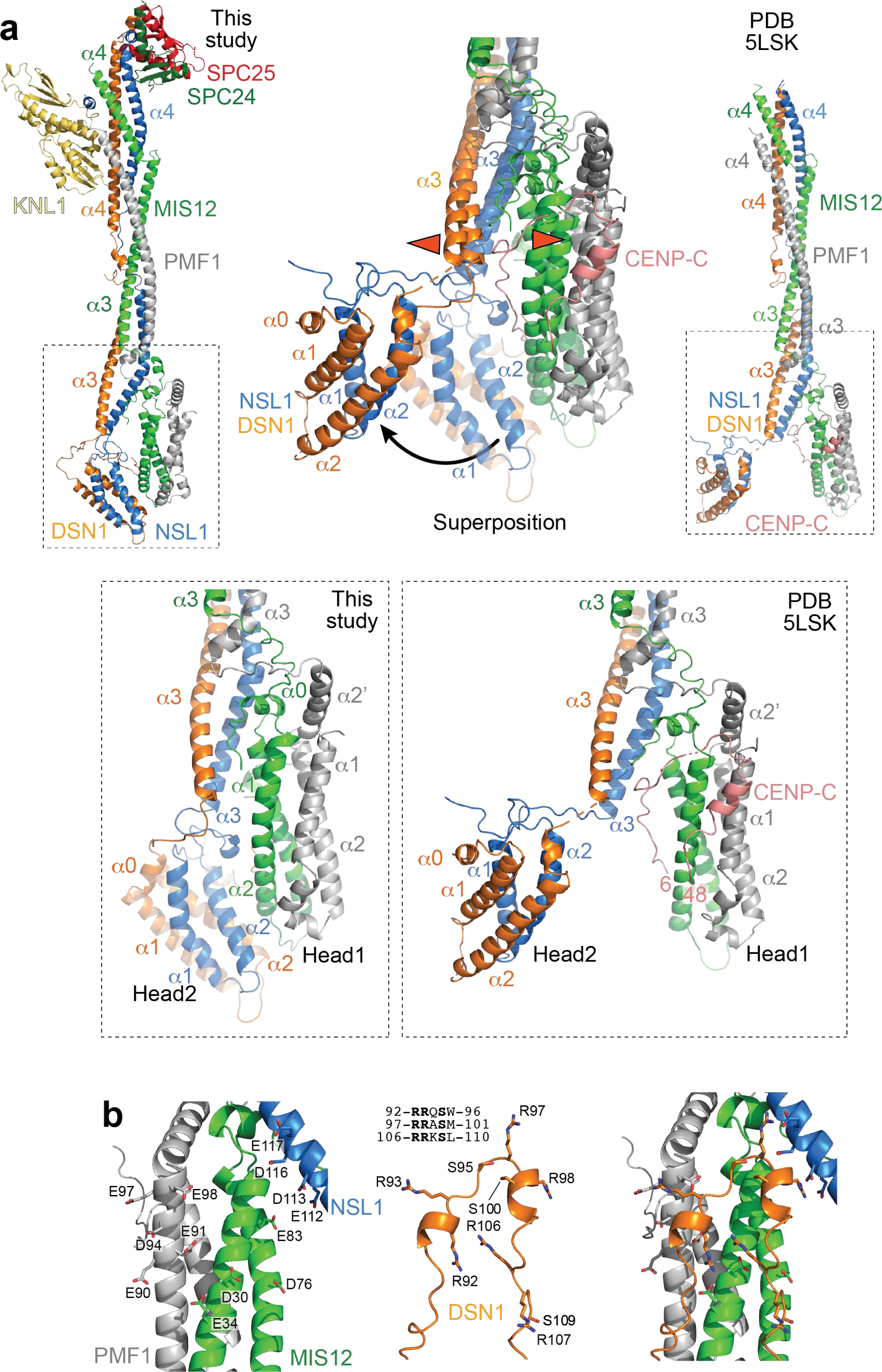
Additional data supporting. **Figure 1** (**a**) Comparison of the closed conformation of the heads observed in this study and the open conformation of the heads observed in a previous structure of the complex of Mis12C with CENP-C (PDB 5LSK; ^22^). In the central superposition, the movement that sways the head2 out of position is marked by a black arrow. The red arrowheads indicate an outward movement of the helices lining the cleft that separates the DSN1 and MIS12 helices when CENP-C binds. (**b**) AF2 model of the interface between a positively-charged regulatory loop of DSN1 and the negatively charged head1 domain (with contributions from NSL1 and DSN1 in head2).

**Figure 2 – Supplement 1.**
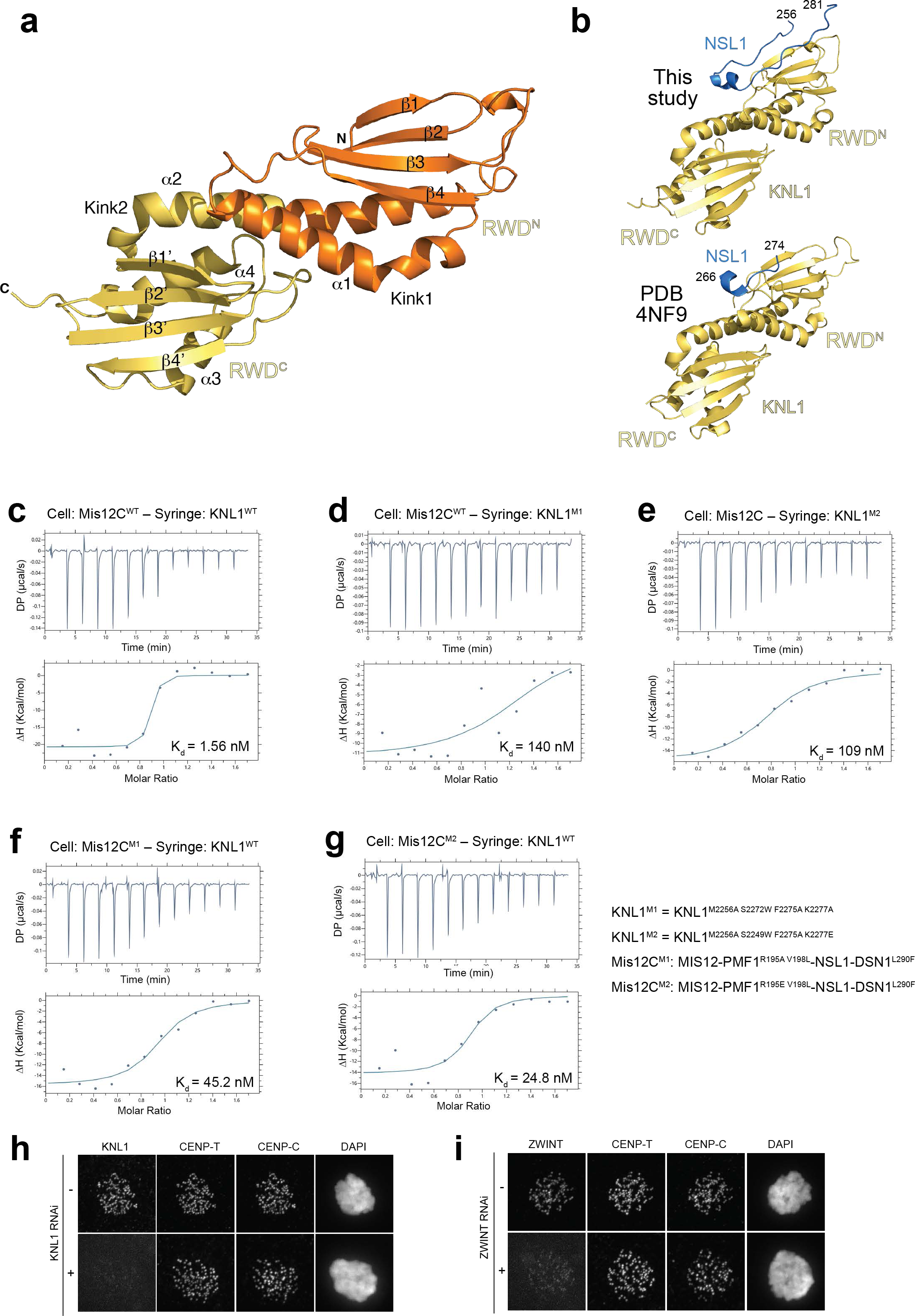
Additional data supporting. **Figure 2** (**a**) Definition of secondary structure of the two RWD domains of the KNL1 C-terminal region, as originally defined in reference ^29^. (**b**) Comparison of the interaction of the NSL1 C-terminal tail with KNL1 as emerging from our current and previous work ^29^. The model of the NSL1 chain in this representation includes extensions, built by AlphaFold2, to the helical segment shown in Figure 2a. Details in Methods and main text. (**c**-**g**) ITC thermograms for the experiments tabulated in Figure 2c. Where applicable, the mutated residues are shown. (**h**) Demonstration of efficiency of protocol of RNAi-based depletion of KNL1. (**i**) Demonstration of efficiency of protocol of RNAi-based depletion of ZWINT.

**Figure 3 – Supplement 1.**
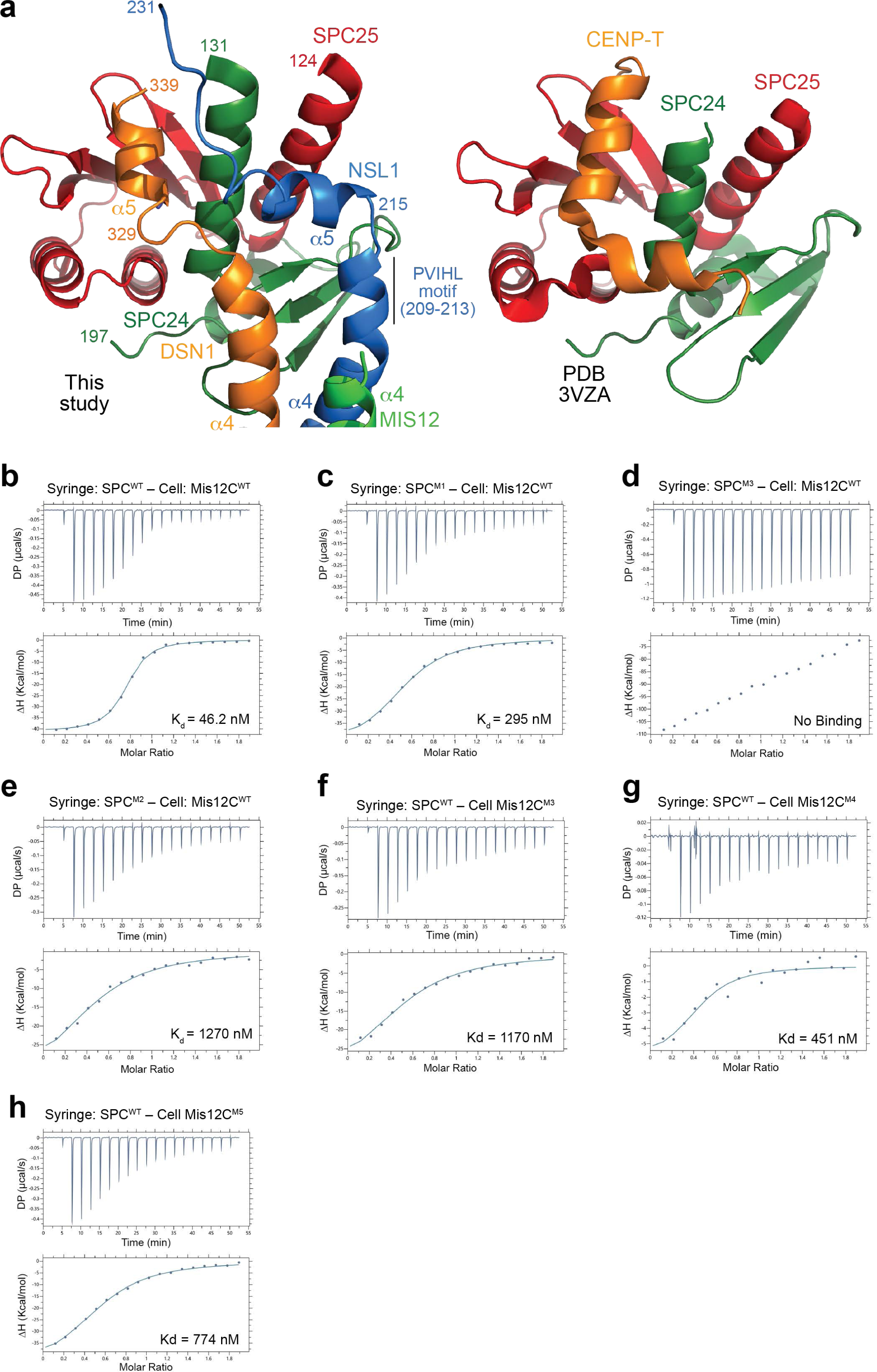
Additional data supporting. **Figure 3** (**a**) Cartoon models of the interaction of the human (left, this study) and avian (PDB ID 3VZA) SPC24/SPC25 complexes, displayed in the same orientation. CENP-T is shown in orange to emphasize the structural similarity with DSN1. (**b**-**h**) Thermograms for the experiments tabulated in Figure 3c.

**Figure 4-S1.**
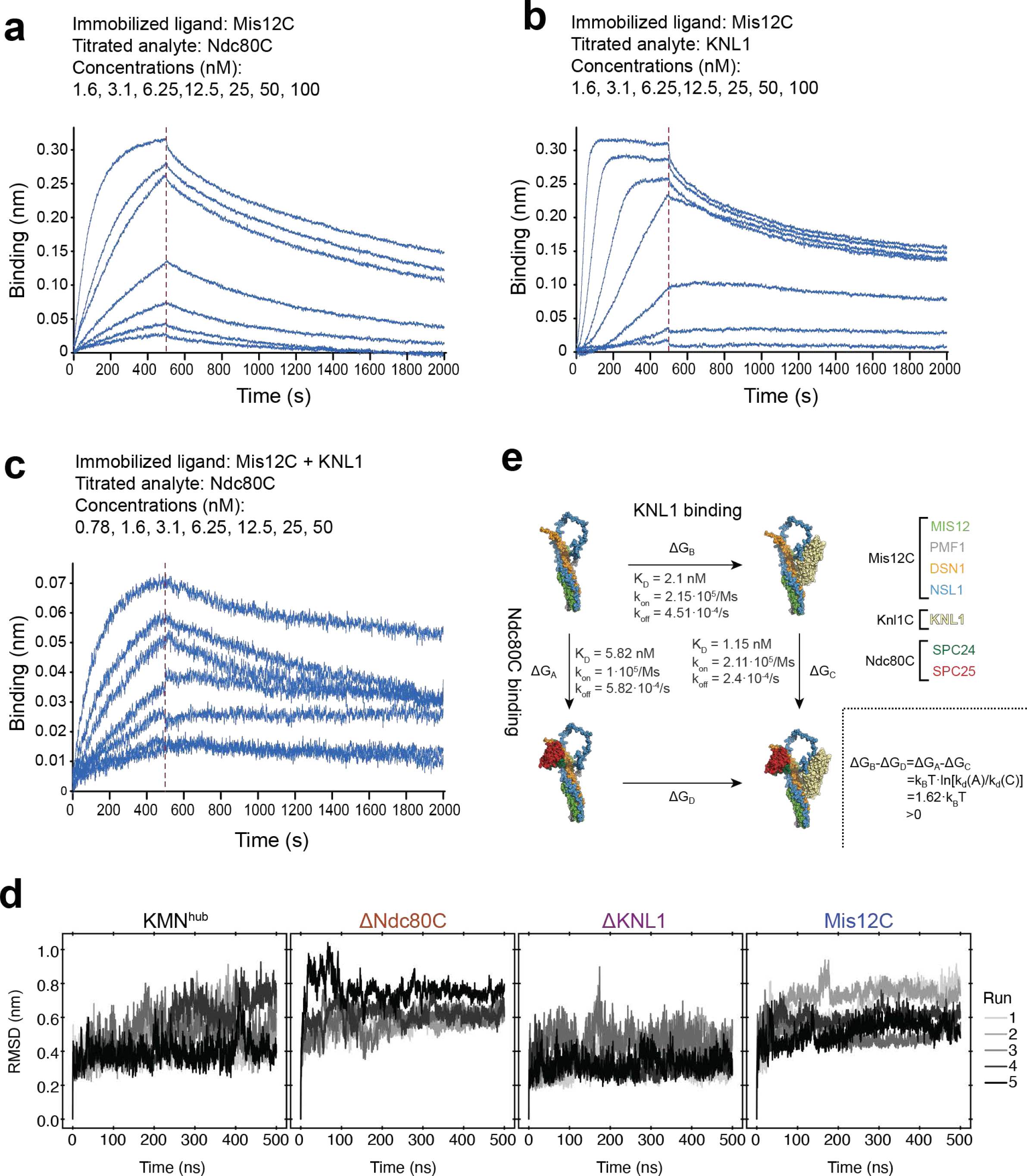
Additional data supporting. **Figure 4** (**a**-**c**) BLI experiments on the indicated species. (**d**) Root mean square deviation (RMSD) from the initial conformation recovered from molecular dynamics simulations of the four indicated systems. Each line represents one independent simulation replica (n=5). (**e**) The thermodynamic cycle depicts the binding of KNL1 (horizontal arrows) or Ndc80C (vertical arrows) to Mis12C, in the absence (top and left arrows) or in the presence (down and right arrows) of the other binding partner. Each possible state shows the protein in sphere representation (colour of protein domains indicated at the right). The measured K_d_, k_on_, and k_off_ values for each respective transition are indicated, together with the respective free energy ΔG. The change related to the binding of Knl1 (ΔGB-ΔGD) was inferred from the affinities to Ndc80C (lower-right box). This change was larger than zero, by almost twice the thermal energy (k_B_T, with k_B_ being the Boltzmann constant and T the temperature). This implies that Ndc80C thermodynamically stabilizes the Mis12C/KNL1 complex, as KNL1 reciprocally does on the Mis12C/Ndc80C interaction.

**Figure 5 – Supplement 1.**
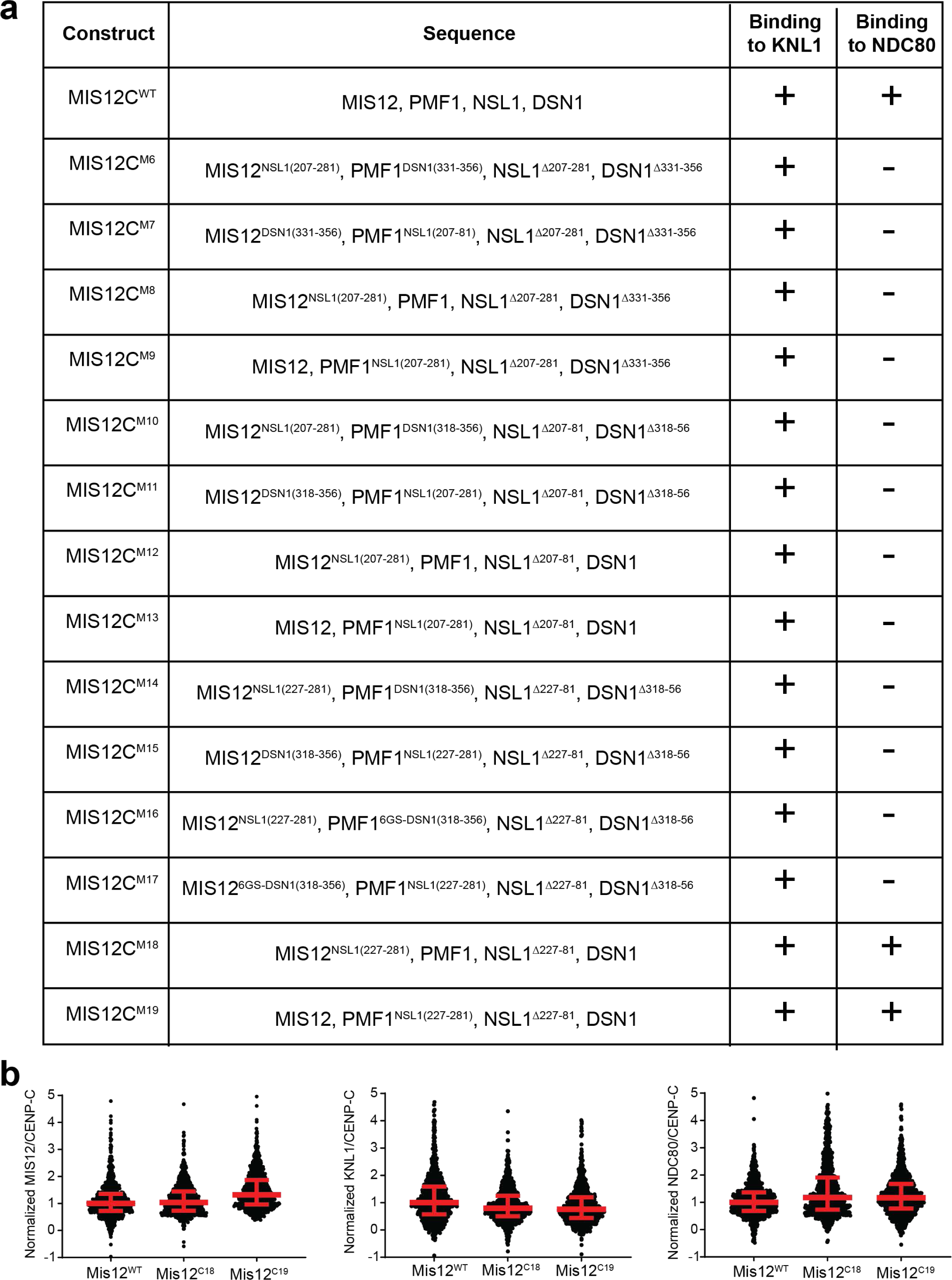
Additional data supporting. **Figure 5** (**a**) Table of swap constructs discussed in the main text. Binding proficiency (+) or lack thereof (-) was evaluated by size-exclusion chromatography as shown for the two constructs in Figure 5b-c. (**b**) Further quantifications for the experiment in Figure 5d.

**Figure 6 – Supplement 1.**
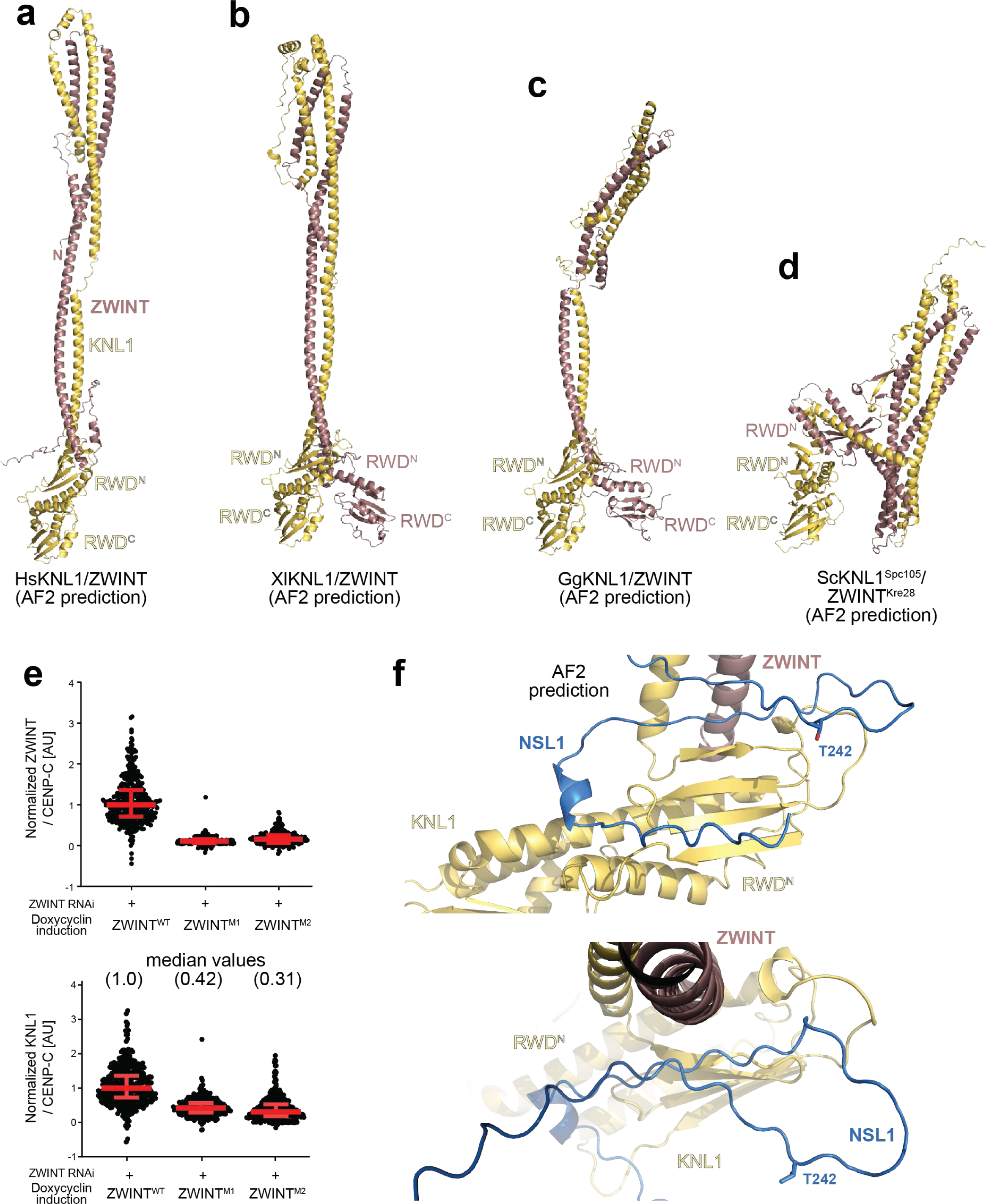
Additional data supporting. **Figure 6** (**a**-**d**) AF2 prediction of KNL1/ZWINT complexes in the indicated species, including a repetition of the human complex in panel **a**. (**e**) Quantification of the experiments in Figure 6g. Red bars represent median and interquartile range of normalized kinetochore intensity values from ZWINT (n = 391), Z1 (n = 291), Z2 (n = 308) from three independent experiments. (**f**) Cartoon model for an AF2 prediction of the NSL1 C-terminal tail bound to the KNL1/ZWINT complex. The position of a phosphorylated residue potentially involved in KNL1 regulation is indicated (see main text for details).

**Figure 7 – Supplement 1.**
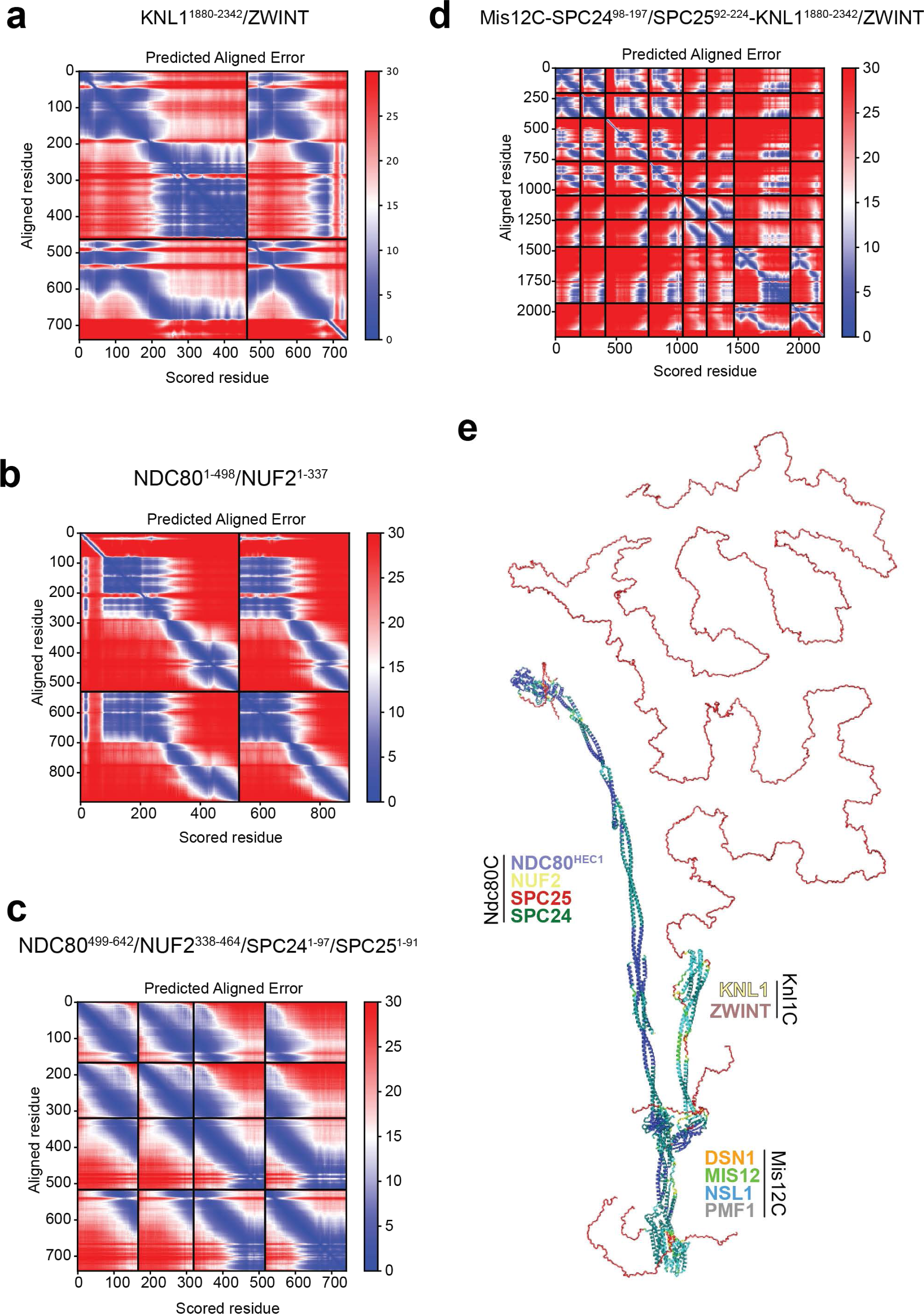
AF2 predictions. (**a**-**d**) AF2 predicted aligned error (PAE) plots for predictions of the indicated constructs. The human KNL1/ZWINT prediction is depicted in Figure 6a. (**e**) Per-residue confidence scores (pLDDT) mapped on the model displayed in Figure 7.

**Table S1.** Cryo-EM data collection, refinement and validation statistics.

|  | KMNZ complex<br>(EMDB-xxxx)<br>(PDB xxxx) |
| --- | --- |
| **Data collection and processing** |  |
| Magnification | 105,000 |
| Voltage (kV) | 300 |
| Electron exposure (e-/Å<sup>2</sup>) | 63.9 |
| Defocus range (μm) | -1.5 to -3.0 |
| Pixel size (Å) | 0.68 (physical) |
| Symmetry imposed | C1 |
| Initial particle images (no.) | 1,792,983 |
| Final particle images (no.) | 230,597 |
| Map resolution (Å) | 4.50 |
| FSC threshold | 0.143 |
| Map resolution range (Å) |  |
| **Refinement** |  |
| Initial model used (PDB code) | AF2 model |
| Model resolution (Å) | 5.5 |
| FSC threshold | 0.5 |
| Map sharpening *B* factor (Å<sup>2</sup>) | DeepEMhancer |
| Model composition |  |
| Non-hydrogen atoms | 9,575 |
| Protein residues | 1,173 |
| Ligands | - |
| *B* factors (Å<sup>2</sup>) |  |
| Protein | 101.0 |
| Ligand | - |
| R.m.s. deviations |  |
| Bond lengths (Å) | 0.004 |
| Bond angles (°) | 0.815 |
| Validation |  |
| MolProbity score | 1.92 |
| Clashscore | 15.6 |
| Poor rotamers (%) | 0.0 |
| Ramachandran plot |  |
| Favored (%) | 96.5 |
| Allowed (%) | 3.4 |
| Disallowed (%) | 0.1 |

**Movie S1 *Molecular dynamics simulations of the four simulated systems***

